# GPCR kinase 3 phosphorylates atypical chemokine receptor 4 independent of G proteins

**DOI:** 10.64898/2026.01.09.698634

**Authors:** Thomas D. Lamme, Isabel B. Sánchez Arroyo, Martine J. Smit, Christopher T. Schafer

## Abstract

Atypical chemokine receptors (ACKRs) indirectly mediate cell migration through chemokine scavenging, which generally requires phosphorylation by G protein-coupled receptor (GPCR) kinases (GRKs) to efficiently control chemokine levels. Despite not coupling G proteins, ACKR4 is preferentially modified by GRK3, a kinase dependent on active G protein subunits for membrane translocation and phosphorylation activity. Here we resolve the underlying mechanisms allowing ACKR4 to circumvent the G protein requirement for GRK3 function. Using live cell BRET assays, we confirm that ACKR4 is preferentially phosphorylated by the GRK2/3 kinase family and that both GRK recruitment and receptor phosphorylation occur in the absence of activated G proteins. Instead, the kinases are recruited directly by a unique acidic rich motif in the proximal receptor C-terminus which coordinates productive phosphorylation reactions. Mutations in this region severely attenuated kinase recruitment and phosphorylation. Productive phosphorylation reaction plays a substantial role in the G protein-independent mechanism and a ‘kinase-dead’ GRK3 (KD-GRK3) has severely reduced recruitment to ACKR4. This was not observed for KD-GRK3 translocation to GPCRs that recruit the kinase in a G protein-dependent manner. Together, these findings suggest that ACKR4 directly coordinates GRK3 recruitment and phosphorylation, highlighting a uniquely evolved atypical mechanism to utilize GRK2/3 while bypassing G protein activation and thereby supporting efficient chemokine scavenging by the atypical receptor.

## Introduction

Chemokine receptors are class A G protein-coupled receptors (GPCRs) that regulate cell migration and contribute to immune homeostasis, inflammation, and development ^1^. Canonical chemokine receptors (CCKRs) activate G protein signaling pathways which drive cell movement along localized chemokine gradients towards higher chemokine concentrations. These gradients are generated and maintained by specialized atypical chemokine receptors (ACKRs) which regulate agonist availability through chemokine scavenging ^2,3^. Unlike CCKRs, ACKRs do not couple G proteins, yet receptor activation still induces GRK-mediated phosphorylation and arrestin recruitment. As a result, ACKRs are often considered β-arrestin-biased ^4^.

ACKR4, previously CCR11 ^5^, CCRL1 ^6^, and CCX-CKR ^7^, regulates chemokine levels by scavenging the chemokines CCL19, CCL20, CCL21, CCL22, and CCL25, thereby limiting ligand availability for CCKRs CCR4, CCR6, CCR7, and CCR9, respectively ^8–11^. ACKR4 is primarily expressed in lymphatic and thymic endothelial cells ^12,13^, which are located adjacent to CCR7- and CCR9-expressing cells such as dendritic cells, T cells, and thymocytes ^14–16^. Chemokine scavenging by ACKR4 facilitates the migration of immune cells by crafting gradients of chemokines shared with the canonical receptors. Like other ACKRs, ACKR4 does not activate heterotrimeric G proteins and chemokine binding leads to phosphorylation by primarily GRK3 ^17^. Arrestins are recruited by the phosphorylated receptor which supports, but is not essential for, chemokine uptake.

Phosphorylation of GPCRs is coordinated by seven kinases (GRK1-7). GRK1 and GRK7 are expressed solely in the retina, while GRK4 expression is restricted to the testes and brain ^18^. The remaining kinases, GRK2, GRK3, GRK5, and GRK6, are ubiquitously expressed, but differ in their subcellular localization. GRK5 and GRK6 are constitutively membrane-associated and anchored by basic residue-lipid interactions or palmitoylation, respectively ^19,20^. GRK2 and GRK3, in contrast, are cytosolic and are recruited to the membrane via complexing with lipidated G⍺_q_ and Gβγ subunits liberated following heterotrimer activation ^21^. Modification of GPCRs by specific kinases is reported to produce distinct phosphorylation patterns ^22^. Therefore, the expression profile of different GRKs can result in cell type specific responses and receptor regulation ^23^.

GPCRs are preferentially phosphorylated by different GRKs and can be generally divided into three main categories: GRK2/3-, GRK5/6-, or GRK2/3/5/6-dependent ^21^. Receptors that couple G proteins, such as CCKRs, are GRK2/3-dependent or GRK2/3/5/6-dependent, reflecting the importance of GRK2/3 phosphorylation when G protein activation is present. β-Arrestin-biased receptors, such as ACKRs, typically rely on GRK5/6 phosphorylation, since these proteins are unable to activate G proteins themselves and thus are unable to recruit GRK2/3 ^23,24^. For example, ACKR3 is dominantly phosphorylated by GRK5/6 ^25,26^, but can borrow Gβγ from co-activated CXCR4 which facilitates GRK2/3 phosphorylation of the atypical receptor in some cellular contexts ^26^. Recently GRK3 and to a lesser extent GRK2 were identified as the primary kinases recruited to ACKR4 ^17^. Given that GRK2/3 phosphorylation is typically G protein-dependent, ACKR4’s preference for GRK2/3 is unexpected.

Here we show that ACKR4 does not require G protein coordination for GRK2/3 recruitment or phosphorylation. Instead, an acidic rich motif in the C-terminus of ACKR4 directly facilitates GRK2/3 recruitment and activity. Recruitment of the kinases to ACKR4 is nearly eliminated when the kinase activity is lost, which is not the case for G protein-dependent GPCRs. These findings suggests that the acidic C-terminus directly coordinates GRK2/3 recruitment by serving as a substrate for modification, representing a novel mechanism by which ACKRs can circumvent canonical GRK dependency on G protein activation.

## Materials and Methods

### Materials

All chemicals and reagents were obtained from Melford or Sigma-Aldrich, unless otherwise specified. HEK293 ΔGRK2/3, HEK293 ΔGRK2/3, HEK293 ΔQ, and corresponding parental HEK293 cell lines were kindly gifted from Carsten Hoffmann (Friedrich-Schiller-Universität Jena) ^27^.

### DNA constructs and site-directed mutagenesis

Human ACKR4 (1-350), FLAG-ACKR3 (2-362), CXCR4 (1-352), and CCR9 (1-369) were inserted into a pcDNA3.1 expression vector, either alone or C-terminally fused to *Renilla* luciferase II (RlucII). β-Arrestin2-GFP10 (kindly provided by N. Heveker, Université de Montréal), GRK3-CT ^26^ (Bovine, residues 547-688, kindly provided by N. Lambert, Augusta University), Gβ_1_ and Gγ_2_ ^28^ (kindly provided by A. Inoue, Tohoku University), GRK3-Nluc ^17^ (kindly provided by D. Legler, Biotechnology Institute Thurgau), mV-CAAX ^29^ (kindly provided by M. Bouvier, Université de Montréal), GRK3 and GRK3_K220R ^27^ (kindly provided by C. Hoffmann, Friedrich-Schiller-Universität Jena) were described previously. FLAG-ACKR3(2-362)_CT(ACKR4)(314-350) followed by a C-terminal RlucII was ordered from Genscript. ACKR4 DE/A, ACKR4 ST/A, GRK3-Nluc RK/A mutants were generated using the Q5 Site-Directed Mutagenesis Kit (New England Biolabs) and validated by Sanger sequencing.

### Chemokine purification from E. coli

Expression and purification of CCL25 and CXCL12 in *E.coli* were performed as previously described ^30,31^. In short, chemokine sequences were inserted into a pET21 vector N-terminally fused to 8His-tag and enterokinase cleavage site. The synthesized plasmid was introduced into the BL21(DE3)pLysS cells, and expression was induced by IPTG. Chemokine-containing inclusion bodies were isolated via sonication and solubilized in a buffer consisting of 50 mM Tris, 6 M guanidine-HCl, and 300 mM NaCl at pH 8.0. Purification was done using a nickel-nitrilotriacetic acid (Ni-NTA) column, washed with 50 mM Mes, 6 M guanidine–HCl, 300 mM NaCl, pH 6.0 and followed by elution with 50 mM acetate, 6 M guanidine–HCl, 300 mM NaCl (pH 4.0). The purified chemokine was reduced using 4 mM DTT and subsequently refolded in 50 mM Tris, 700 mM arginine-HCl, 1 mM EDTA, 200 mM glutamine, 0.1% Triton-X and 1 mM GSSG at pH 7.5 by dropwise addition. After refolding, the chemokine was dialyzed in 20 mM Tris (pH 8.0) with 150 mM NaCl. The 8His-tag was cleaved of using Enterokinase (New England Biolabs) and verified by SDS-PAGE and LC-MS analysis. The cleaved product was further refined on a Ni-NTA column, with washes in 50 mM Tris (pH 8.0) and elution in 6 M guanidine-HCl with 50 mM MES (pH 6.0) or 50 mM acetate (pH 4.0) as two separate fractions. The fractions were purified by reverse-phase HPLC using a Gemini C18 110A column (Phenomex), using a linear gradient of 5-95% acetonitrile in 0.1% TFA. The purified chemokines were collected, lyophilized and stored at -80 °C.

### Transfection HEK293 cells in suspension

HEK293 were maintained in Dulbecco’s modified Eagle’s medium (Thermo Fisher Scientific) with 10% fetal bovine serum (Bodinco) and 1% penicillin and streptomycin (Gibco) and incubated at 37 °C in a humidified 5% CO_2_ environment. Transfection was performed with a total of 2 μg plasmid DNA per 1 x 10^6^ cells using 6 μg polyethyleneimine (PEI; Polysciences, inc.) in 150 mM NaCl. DNA-PEI complexes were allowed to form by incubation for 15 min at room temperature. In the meantime, cells were harvested with trypsin-EDTA (Thermo Fisher Scientific) and counted before addition to the DNA-PEI complexes. Cells were plated at 30k/well in a white 96-well plate (Greiner) and incubated for 48 h. Routine testing confirmed the absence of mycoplasma contamination.

### β-arrestin2 recruitment by BRET

HEK293, including ΔGRK2/3, ΔGRK5/6, and ΔQ knockout lines, were transiently transfected with 50 ng of ACKR4-RlucII/ACKR3-RlucII/CXCR4-RlucII/CCR9-RlucII/ACKR3_CT4-RlucII and 1 µg of GFP10-β-arrestin2. If indicated, cells were co-transfected with either 950 ng of GRK3-CT, 450 ng of Gβ_1_ and Gγ_2,_ or 100 ng of GRK2/GRK2 R587Q/GRK3/GRK3 R587Q. Total of 2 µg DNA per condition was reached by supplementing with empty pcDNA3.1 plasmid. Transfections were carried out using standard methods described previously. After 48 h, cells were gently rinsed once with PBS and maintained in Hank’s Balanced Salt Solution (HBSS) supplemented with 0.1% Bovine Serum Albumin (BSA; Fraction V; PanReac AppliChem). Next, cells were incubated with 5 µM Prolume Purple (Prolume Ltd) for 5 min at room temperature to allow substrate diffusion and stabilization of luminescence. Three baseline readings were acquired prior to stimulation. Subsequently, cells were stimulated with increasing concentrations of CCL25 or CXCL12. Bioluminescence was quantified using a PHERAstar plate reader equipped with a dual emission filter (410-80 nm and 515-30 nm) for 40 min at 37 °C. BRET ratio was calculated by the intensity at 515-30 nm, divided by the intensity at 410-80 nm. Results are from three independent experiments reported as AUC of the kinetic ΔBRET trace (as shown in **Fig. 1b**) normalized to WT response. The AUC was selected as a measure since the kinetic profiles (over 40 min) of the various chemokine receptors differed, making a comparison at a single time point potentially misleading.

**Figure 1.**
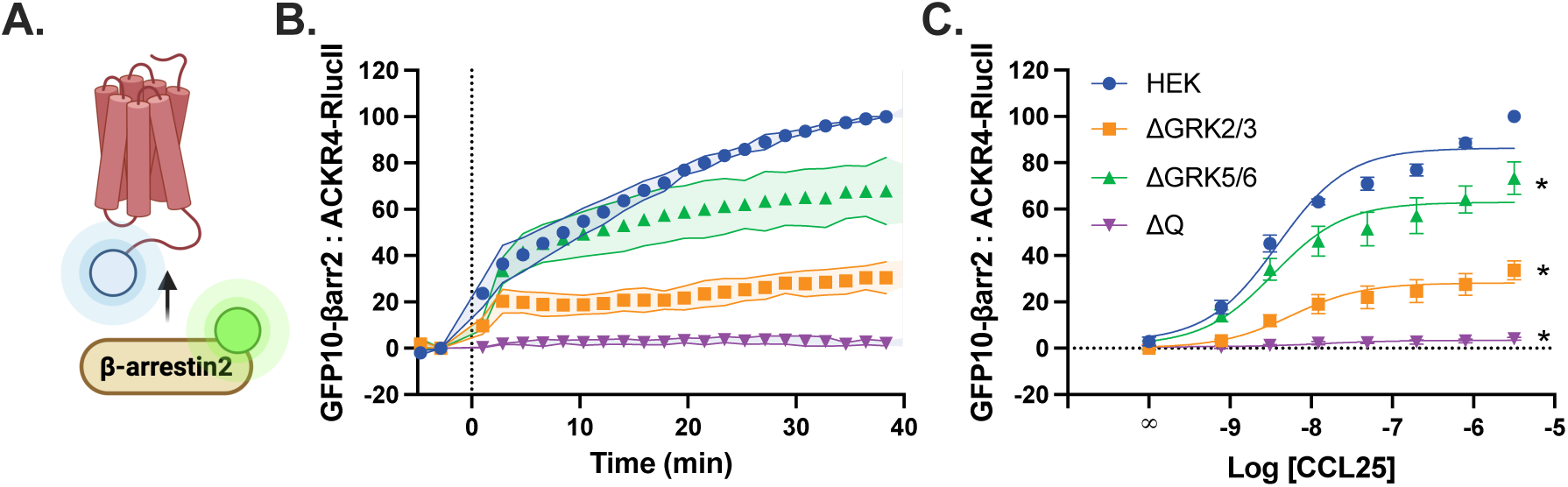
β-Arrestin2 recruitment to ACKR4 is primarily mediated by GRK2/3 phosphorylation. (A) Schematic illustration of BRET pairs ACKR4-RlucII and GFP10-βarr2. βarr2 recruitment following stimulation with 100 nM CCL25 over time (B) or across a titration of CCL25 concentrations (C) towards ACKR4. Values represent the mean ± SD of three independent experiments performed in triplicate, normalized to maximal HEK response. Statistical significance at the top of the sigmoidal curve was determined by using the extra sum-of-squares F test ∗p < 0.0001.

### Bystander GRK3 recruitment by BRET

HEK293 cells were transiently transfected with 100 ng ACKR4/FLAG-ACKR3/CXCR4/CCR9, 50 ng of GRK3-Nluc, and 200 ng of mV-CAAX. If indicated, cells were co-transfected with 950 ng GRK3-CT. Total of 2 µg DNA per condition was reached by supplementing with empty pcDNA3.1 plasmid. Transfections were carried out using standard methods described previously. After 48 h, cells were gently rinsed once with PBS and maintained in HBSS supplemented with 0.1% BSA. Next, cells were incubated with 2.5 µM Furimazine (Promega) for 5 min at room temperature to allow substrate diffusion and stabilization of luminescence. Three baseline readings were acquired prior to stimulation. Subsequently, cells were stimulated with 100 nM of CCL25 or CXCL12. Bioluminescence was quantified using a PHERAstar plate reader equipped with a dual emission filter (475-30 nm and 535-30 nm) for 40 min at 37 °C. BRET ratio was calculated by the intensity at 535-30 nm, divided by the intensity at 475-30 nm. Results are from three independent experiments reported as ΔBRET trace. AUC of this kinetic trace was used for comparison between conditions.

### CCL25 uptake by flow cytometry

HEK293 cells were transiently transfected with 100 ng ACKR4 to a total of 2 µg DNA per condition by supplementing with empty pcDNA3.1 following previously described procedures. After 48 h, cells were harvested using Accutase (Thermo Fisher Scientific) and transferred to a Guava-compatible 96-well conical plate (Greiner). Between each step, cells were centrifuged at 350 x g for 3 min and supernatant was decanted. Cells were washed twice with FACS buffer (PBS supplemented with 0.5% BSA). Next, cells were incubated with 100 nM CCL25-AZ488 (Protein Foundry) for 1.5 h at 37 °C while gently shaking. After incubation, cells were washed once with FACS buffer, twice with acidic citrate buffer (50 mM citric acid, 150 mM NaCl, pH 4.5) to wash of residual surface-bound and non-internalized chemokine. Following three washes with FACS buffer, cells were resuspended in the same buffer before analysis on a Guava easyCyte flow cytometer (Cytek). Mean green fluorescence intensities were recorded for at least 5000 cells and normalized to WT and pcDNA3.1 control levels.

### Surface expression by flow cytometry

HEK293 cells were transiently transfected with 100 ng ACKR4 to a total of 2 µg DNA per condition by supplementing with empty pcDNA3.1 following previously described procedures. After 48 h, cells were harvested using Accutase (Thermo Fisher Scientific) and transferred to a Guava-compatible 96-well conical plate (Greiner). All subsequent steps were performed at 4°C and centrifuged at 350 x g for 3 min between washing or incubation step. Cells were washed twice with filtered FACS buffer and incubated with anti-hACKR4 mouse antibody (1:500 dilution) (Biolegend; MAB-13E11) for 1 h. After washing, samples were stained with anti-mouse PE-conjugated secondary antibody (R&D Systems; F0102B) for 1 h in the dark. Next, cells were washed three times before being resuspended in FACS buffer. Mean yellow fluorescence intensities were recorded and normalized to WT and pcDNA3.1 control levels.

### Alphafold3 model building

Structural models of the CCL25:ACKR4:GRK3 complex were predicted using AlphaFold3.0 with default parameters ^32^ and the following amino acid sequences.

CCL25:

QGVFEDCCLAYHYPIGWAVLRRAWTYRIQEVSGSCNLPAAIFYLPKRHRKVCGNPKSREVQRAMKLLDARNKVFAKLHHNTQTFQAGPHAVKKLSSGNSKLSSSKFSNPISSSKRNVSLLISANSGL

ACKR4:

MALEQNQSTDYYYEENEMNGTYDYSQYELICIKEDVREFAKVFLPVFLTIVFVIGLAGNSMVVAIYAYYKKQRTKTDVYILNLAVADLLLLFTLPFWAVNAVHGWVLGKIMCKITSALYTLNFVSGMQFLACISIDRYVAVTKVPSQSGVGKPCWIICFCVWMAAILLSIPQLVFYTVNDNARCIPIFPRYLGTSMKALIQMLEICIGFVVPFLIMGVCYFITARTLMKMPNIKISRPLKVLLTVVIVFIVTQLPYNIVKFCRAIDIIYSLITSCNMSKRMDIAIQVTESIALFHSCLNPILYVFMGASFKNYVMKVAKKYGSWRRQRQSVEEFPFDSEGPTEPTSTFSI

GRK3:

MADLEAVLADVSYLMAMEKSKATPAARASKRIVLPEPSIRSVMQKYLAERNEITFDKIFNQKIGFLLFKDFCLNEINEAVPQVKFYEEIKEYEKLDNEEDRLCRSRQIYDAYIMKELLSCSHPFSKQAVEHVQSHLSKKQVTSTLFQPYIEEICESLRGDIFQKFMESDKFTRFCQWKNVELNIHLTMNEFSVHRIIGRGGFGEVYGCRKADTGKMYAMKCLDKKRIKMKQGETLALNERIMLSLVSTGDCPFIVCMTYAFHTPDKLCFILDLMNGGDLHYHLSQHGVFSEKEMRFYATEIILGLEHMHNRFVVYRDLKPANILLDEHGHARISDLGLACDFSKKKPHASVGTHGYMAPEVLQKGTAYDSSADWFSLGCMLFKLLRGHSPFRQHKTKDKHEIDRMTLTVNVELPDTFSPELKSLLEGLLQRDVSKRLGCHGGGSQEVKEHSFFKGVDWQHVYLQKYPPPLIPPRGEVNAADAFDIGSFDEEDTKGIKLLDCDQELYKNFPLVISERWQQEVTETVYEAVNADTDKIEARKRAKNKQLGHEEDYALGKDCIMHGYMLKLGNPFLTQWQRRYFYLFPNRLEWRGEGESRQNLLTMEQILSVEETQIKDKKCILFRIKGGKQFVLQCESDPEFVQWKKELNETFKEAQRLLRRAPKFLNKPRSGTVELPKPSLCHRNSNGL

Models were analyzed in ChimeraX-1.6.1, including generation of pLDDT plots. PAE plots were generated and analyzed using PAE viewer ^33^.

### Statistical analysis

All statistical analysis were conducted using GraphPad Prism 10. Data representation in figures, including bar plots, symbols, and error bars are specified in the corresponding figure legends. For scatter plots, the values are the mean from three independent experiments, with each experiment performed in triplicate. For bar charts, the bars indicate the mean of the three independent experiments, while the individual points represent the means of each experiment measured in triplicate. All error bars correspond to the standard deviation (SD). For kinetic traces, the SD is depicted as the colored area around the data points. Dose-response curves were fitted using a three-parameter sigmoidal model (log[agonist] versus response) in GraphPad Prism 10, described by the equation below:

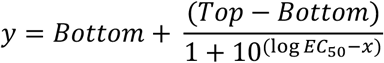

For the AUC analysis, the baseline was set at 0. Statistical significance for the DRCs was assessed by using the extra sum-of-squares F test using the top value for comparison. All other comparisons were done by one-way Brown-Forsythe and Welch ANOVA, followed by a Dunnet’s T3 post hoc test.

## Results

### Arrestin recruitment to ACKR4 is primarily dependent on GRK2/3 phosphorylation

While ACKR4 recruits GRK2/3, it is not clear yet how the four ubiquitously expressed GRKs contribute to receptor phosphorylation. To assess the relative importance of the different GRKs, we monitored Bioluminescence Resonance Energy Transfer (BRET) between a C-terminally-tagged ACKR4 (ACKR4-RlucII) and GFP10-tagged β-arrestin2 (GFP10-βarr2) (**Fig. 1A**) in cells where GRK2/3 (ΔGRK2/3), GRK5/6 (ΔGRK5/6), or GRK2/3/5/6 (ΔQ) were knocked out (KO) by CRISPR ^27^. Arrestins are recruited to the receptor following C-terminal phosphorylation by GRKs and changes to the recruitment reflect altered phosphorylation amounts or patterns. In WT cells, CCL25-stimulation induced a gradually increasing BRET signal, representing an accumulation of arrestin:ACKR4 complexes (**Fig. 1B**). Arrestin recruitment was moderately reduced in GRK5/6 deficient cells, whereas ΔGRK2/3 cells showed a substantially lower response (∼30% of the maximal WT response) (**Fig. 1C, Supplementary Table 1**). In ΔQ cells, where no GRK phosphorylation is possible, no arrestin recruitment was observed with chemokine stimulation. While ACKR4 expression was slightly lower in the KO cells (**Supplemental Fig. 1**), it did not correlate with the relative decreased arrestin engagement. These results indicate that ACKR4 is primarily dependent on GRK2/3 phosphorylation, consistent with previous recruitment observations ^17^. This is particularly unusual for an atypical receptor, since GRK2/3 recruitment to the plasma membrane (PM) is typically G protein-dependent ^23,34,35^ suggesting ACKR4 may be phosphorylated by GRK2/3 via a G protein-independent mechanism.

### GRK2/3 acts on ACKR4 independent of G protein interactions

GRK2/3 localization to the PM is coordinated by Gβγ subunits released from the G protein heterotrimer upon G protein activation and is an essential regulatory step for phosphorylation by these kinases ^34,36,37^. To determine if the GRK2/3 recruitment to ACKR4 still requires Gβγ interactions, we assessed GRK3 translocation to the membrane and phosphorylation (by arrestin interactions) of ACKR4 in the presence of GRK3-CT. GRK3-CT consists of the Gβγ interaction domain of GRK3 and its overexpression inhibits GRK2/3 activity by competing for freed Gβγ (**Fig. 2A**) ^23,37–39^. The effects were compared with receptors known to utilize the canonical G protein-dependent GRK2/3 mechanism, including the chemokine receptors ACKR3 and CXCR4, an atypical and canonical pair sharing the chemokine CXCL12, and the canonical CCL25-binding sibling receptor of ACKR4, CCR9 ^26^. The CCKRs CXCR4 and CCR9 are phosphorylated by both GRK2/3 and GRK5/6 ^30,40^, while ACKR3 is preferentially phosphorylated by GRK5/6 and can be modified by GRK2/3 when supplied with Gβγ from CXCR4 co-activation ^26^. The recruitment of GRK3 to native C-termini receptors (untagged) was tracked by bystander BRET between GRK3-Nanoluciferase (GRK3-Nluc) and mVenus anchored to the PM by a CAAX motif (mV-CAAX). Upon addition of CCL25, ACKR4 shows a quick recruitment of GRK3 towards the membrane, that peaks at ∼5 min and then gradually decreases over time (**Fig. 2B**). Co-expression with GRK3-CT does not significantly affect kinase recruitment by ACKR4, suggesting a Gβγ-independent recruitment. ACKR3 stimulation shows limited recruitment of GRK3 towards the membrane that quickly returns to baseline (**Fig. 2C**), in line with the receptor’s established dependency on GRK5/6 and inability to generate its own Gβγ for GRK3 recruitment ^23,26,41^. The minimal translocation of GRK3 to ACKR3 is eliminated with co-expression of GRK3-CT confirming Gβγ dependency. Both CXCR4 (**Fig. 2D**) and CCR9 (**Fig. 2E**) show robust GRK3 recruitment, which is significantly reduced when co-expressed with GRK3-CT. Compared to the other receptors, GRK3 recruitment to ACKR4 is much slower and may suggest unassisted translocation from the cytosol.

**Figure 2.**
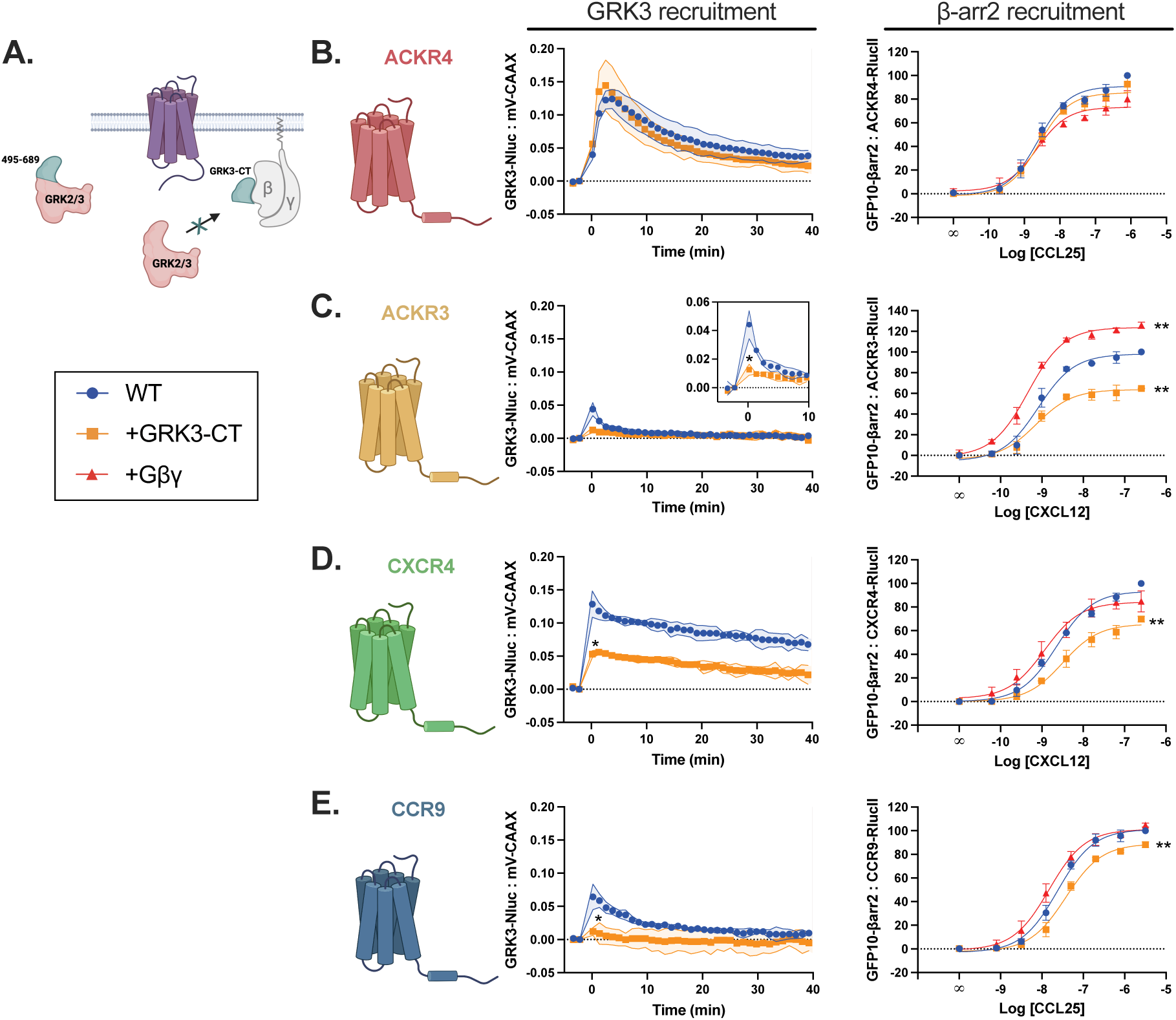
ACKR4 recruits GRK2/3 independent of Gβγ. (A) Schematic illustration of GRK3-CT as an effective competitor for the GRK2/3:Gβγ interaction. GRK3 localization is measured by bystander BRET between GRK3-Nluc and mV-CAAX with or without co-expression of GRK3-CT after stimulation with 100 nM chemokine of untagged ACKR4 (B, left), ACKR3 (C, left), CXCR4 (D, left), or CCR9 (E, left) measured over time. Recruitment of GFP10-βarrestin2 to ACKR4-RlucII (B, right), ACKR3-RlucII (C, right), CXCR4-RlucII (D, right), or CCR9-RlucII (E, right) across a titration of chemokine concentrations with co-expression of GRK3-CT or Gβγ as indicated. Values represent the mean ± SD of three independent experiments performed in triplicate. Statistical significance at the top of the sigmoidal curve was determined by using the extra sum-of-squares F test or by an unpaired t test compared at the peak (GRK3 bystander recruitment). ∗p < 0.05, ∗∗p < 0.0001.

Next, the impact of Gβγ on the receptor phosphorylation state was assessed by BRET-based β-arrestin2 recruitment (**Fig. 1A**). Co-expression of GRK3-CT had no effect on the chemokine-induced arrestin recruitment to ACKR4 (**Fig. 2B**), but showed a significant decrease for ACKR3, CXCR4, and CCR9 (**Fig. 2C, D, and E**), mirroring the impact of GRK3-CT on GRK3 recruitment. We have previously shown the co-expression of the Gβγ subunits can bypass the need for G protein activation for GRK2/3 phosphorylation of atypical GPCRs ^26^. Circumventing the G proteins had no effect on ACKR4 arrestin recruitment (**Fig. 2B**), further supporting a Gβγ-independent mechanism for ACKR4 phosphorylation. In contrast, Gβγ co-expression enhanced arrestin recruitment to ACKR3 (**Fig. 2C**), consistent with previous findings that the receptor relies on Gβγ for GRK2/3 phosphorylation despite not producing its own ^26^. No significant changes were observed for CXCR4 and CCR9 (**Fig. 2D and E**), likely because these receptors produce sufficient Gβγ through G protein activation to achieve maximal arrestin recruitment. Together this shows that ACKR4, but not ACKR3, CXCR4, and CCR9, can recruit GRK3 independent of Gβγ.

To confirm that GRK2/3 indeed do not require G protein interactions to phosphorylate ACKR4, we tested the ability of GRK2 and GRK3 mutants with impaired Gβγ interactions to phosphorylate the panel of receptors. The mutation R587Q (same numbering in both kinases) is located in the pleckstrin homology domain at the Gβγ binding interface and disrupts complex formation (**Fig. 3A**) ^23,41^. WT and mutant GRK2 and GRK3 were expressed in cells lacking endogenous GRKs (ΔQ) and their impact on arrestin recruitment was determined by BRET ^27^. Reintroduction of WT GRK2/3 lead to a marked increase in basal BRET values between ACKR4 and arrestin (**Fig. 3B**) consistent with previously documented constitutive activity ^17^. Basal arrestin interactions in the presence of GRK2 R587Q and GRK3 R587Q expression was still observed, albeit much less than compared to the WT GRKs. Receptor activation by CCL25 resulted in increased arrestin recruitment by GRK2/3 R587Q compared to the WT GRKs, suggesting that the CCL25-induced arrestin recruitment by ACKR4 does not require the GRK2/3:Gβγ interaction (**Fig. 3B**). In fact, blocking this interaction slightly increased the efficacy of chemokine induced arrestin recruitment, which may indicate that more GRK2/3 R587Q are available to phosphorylate ACKR4 as canonical receptors cannot effectively utilize these GRKs. Alternatively, the greater basal arrestin interaction with WT GRK co-expression may limit how large of a change can be promoted by chemokine addition, leading to a greater change in BRET with the mutant GRKs. ACKR3 is constitutively active ^42–44^ and also recruits arrestins constitutively in the presence of WT GRKs, but this interaction is abolished with the Gβγ-deficient mutant kinases (**Fig. 3C**). The CXCL12-induced effect shows impaired arrestin recruitment to ACKR3 for the mutated GRKs compared to WT. No basal arrestin interaction was observed for CXCR4 (**Fig. 3D)** and chemokine-promoted arrestin recruitment was significantly lower for the GRK mutants compared to WT GRK2/3. These results are consistent with ACKR3 and CXCR4 both requiring the GRK2/3:Gβγ interaction for GRK2/3 phosphorylation, while ACKR4 is phosphorylated by an independent mechanism.

**Figure 3.**
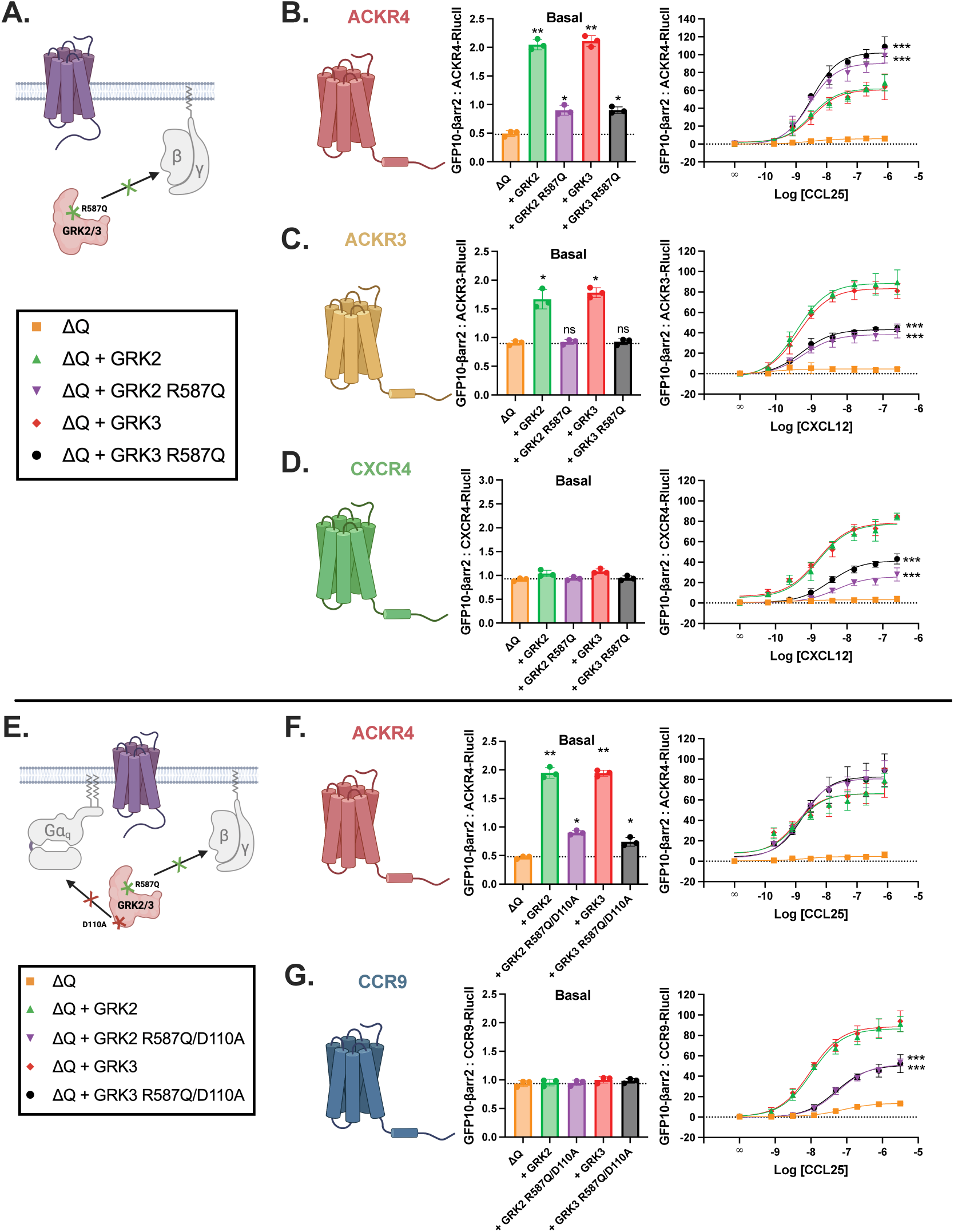
ACKR4 does not need GRK2/3:Gβγ or GRK2/3:G⍺_q_ interactions, unlike ACKR3, CXCR4 and CCR9. (A) Schematic illustration of point mutation R587Q that impairs the interaction of GRK2/3 with Gβγ. Constitutive (left) and chemokine-induced (right) arrestin recruitment measured by BRET between GFP10-βarr2 and ACKR4-RlucII (B), ACKR3-RlucII (C), and CXCR4-RlucII (D) in ΔQ across a titration of chemokine concentrations. Co-expression in ΔQ cells with WT GRK2/3 or GRK2/3 R587Q if indicated. (E) Schematic illustration of double mutant of GRK2/3 that lowers the affinity for both Gβγ and G⍺_q_. Constitutive (left) and chemokine-induced (right) arrestin recruitment measured by BRET between GFP10-βarr2 and ACKR4-RlucII (F) and CCR9-RlucII (G) in ΔQ cells across a titration of chemokine concentrations. Co-expression in ΔQ cells with WT GRK2/3 or GRK2/3 D110A/R587Q if indicated. Values represent the mean ± SD of three independent experiments performed in triplicate. Statistical significance at the top of the sigmoidal curve was determined by using the extra sum-of-squares F test, whereas significance for bar graphs was determined using Welch ANOVA followed by a Dunnett’s T3 multiple comparisons test (basal). ∗p < 0.05, ∗∗p < 0.001, and ∗∗∗p < 0.0001.

The GRK2/3 translocation to the membrane can also be stabilized by interaction with G⍺_q_ ^45^. This interaction can be effectively impaired by a specific point mutation (D110A) in the G protein-signaling homology domain of GRK2/3 ^45^ (**Fig. 3E**). The contribution of G⍺_q_ to GRK2/3 recruitment to ACRK4 was thus assessed by introduction of double mutant (D110A/R587Q) GRKs into ΔQ cells and evaluated by arrestin recruitment. Here, CCR9, a G⍺_i_ and G⍺_q_-coupled CCKR, was taken as a positive control ^30^. Constitutive arrestin recruitment with co-expression of the double GRK mutants showed a similar pattern as with the GRK R587Q mutants for ACKR4 (**Fig. 3F**), while CCL25 induced similar arrestin recruitment with both WT and mutant GRKs. Similar to CXCR4, CCR9 showed no constitutive arrestin interaction (**Fig. 3G**). The CCL25-induced arrestin recruitment to CCR9 was significantly impaired in the mutant GRKs compared to the WT kinases, similar to the effect of R587Q GRKs on ACKR3 and CXCR4. Together these results suggest that phosphorylation of ACKR4 by GRK2/3 is independent of both Gβγ and G⍺_q_ membrane targeting interactions, unlike ACKR3, CXCR4, and CCR9.

### The determinants of Gβγ-independent GRK2/3 phosphorylation are contained in the ACKR4 C-terminus

With canonical GRK2/3 mechanisms coordinated by Gβγ and G⍺_q_ excluded, we hypothesized that ACRK4 may directly facilitate GRK2/3 recruitment to the PM leading to receptor phosphorylation. The receptor C-terminus is a key regulatory domain for most GPCRs and a primary target for GRK phosphorylation. Thus, we predicted that the determinants for direct GRK2/3 interactions may be contained within the ACKR4 C-terminus and these could be transferred to another GPCR. To test this hypothesis, we replaced the C-terminus of ACKR3 with that of ACKR4 following the structurally conserved NPXXY motif at the end of TM7 to preserve all elements of the ACKR4 C-terminal tail (**Fig. 4A**). ACKR3 was chosen as it also does not couple G proteins and thus would predominantly rely on the transferred features of the ACKR4 C-terminus for GRK2/3 recruitment. Arrestin association to the chimeric ACKR3 (ACKR3_CT(ACKR4)) was assessed by BRET with the same addback experiments as presented in Fig. 3. The basal arrestin interaction for the C-terminal substitution is identical to WT ACKR3 with only WT GRKs promoting an increase, suggesting the constitutive activity of ACKR3 is unaffected by domain swap (**Fig. 4B**). Arrestin recruitment promoted by CXCL12 to the chimera mimicked the profile of ACKR4, with an elimination of the differences between the WT and mutant GRKs (**Fig. 4C**). These results suggest that specific features of the ACKR4 C-terminus facilitate non-canonical GRK2/3 recruitment and phosphorylation.

**Figure 4.**
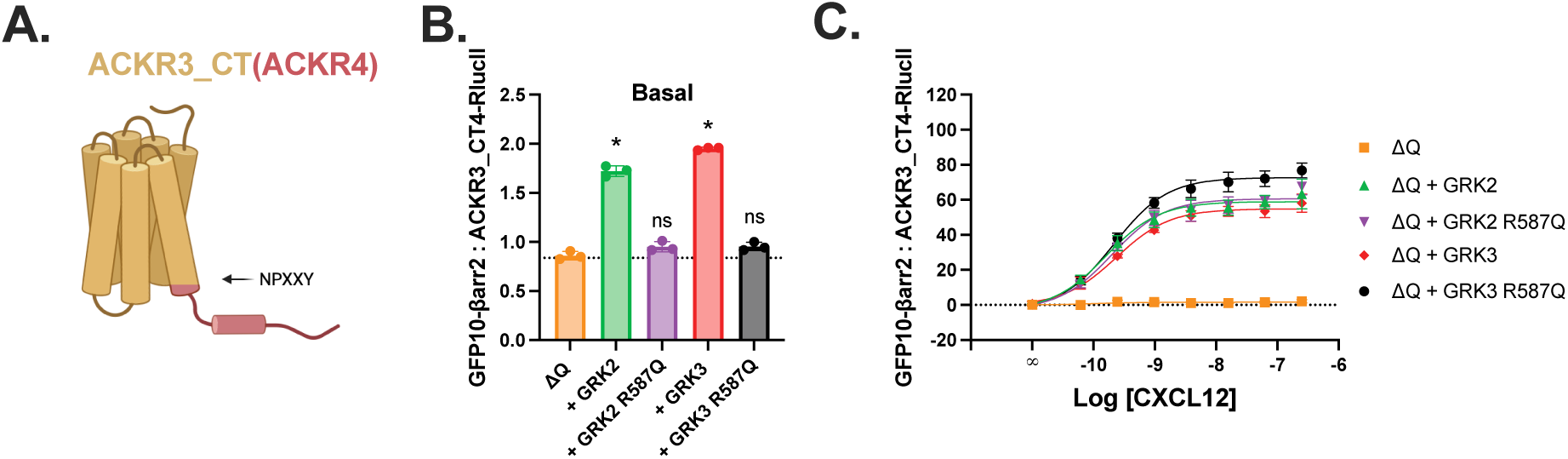
ACKR3-ACKR4 C-terminus chimera makes it independent of Gβγ. (A) Schematic illustration of the chimeric ACKR3 receptor in which the C-terminal tail of ACKR3 was replaced by that of ACKR4 (ACKR3_CT(ACKR4)) after the NPXXY motif. Constitutive (B) and chemokine-induced (C) arrestin recruitment measured as BRET between GFP10-βarr2 and ACKR3_CT(ACKR4)-RlucII in ΔQ cells across a titration of chemokine concentrations. Co-expression in ΔQ cells with WT GRK2/3 or GRK2/3 R587Q as indicated. Values represent the mean ± SD of three independent experiments performed in triplicate. Statistical significance at the top of the sigmoidal curve was determined by using the extra sum-of-squares F test, whereas significance for bar graphs was determined using Welch ANOVA followed by a Dunnett’s T3 multiple comparisons test (basal). ∗p < 0.001

### Unique acidic rich proximal C-terminus is responsible for direct GRK3:ACKR4 interaction

The ACKR3-CT(ACKR4) results (**Fig. 4**) suggest that specific motifs in the ACKR4 C-terminus coordinate G protein-independent GRK2/3 phosphorylation. Therefore, we looked for unique patterns in the C-terminal sequence of ACKR4 compared to other chemokine receptors. We found that ACKR4 has a uniquely acidic-rich proximal C-terminus which features a series of negatively charged amino acids (**EE**XXX**D**X**E**XXX**E**) immediately following the putative end of helix 8 and followed by a high density of potential phosphosites (**Fig. 5**).

**Figure 5.**
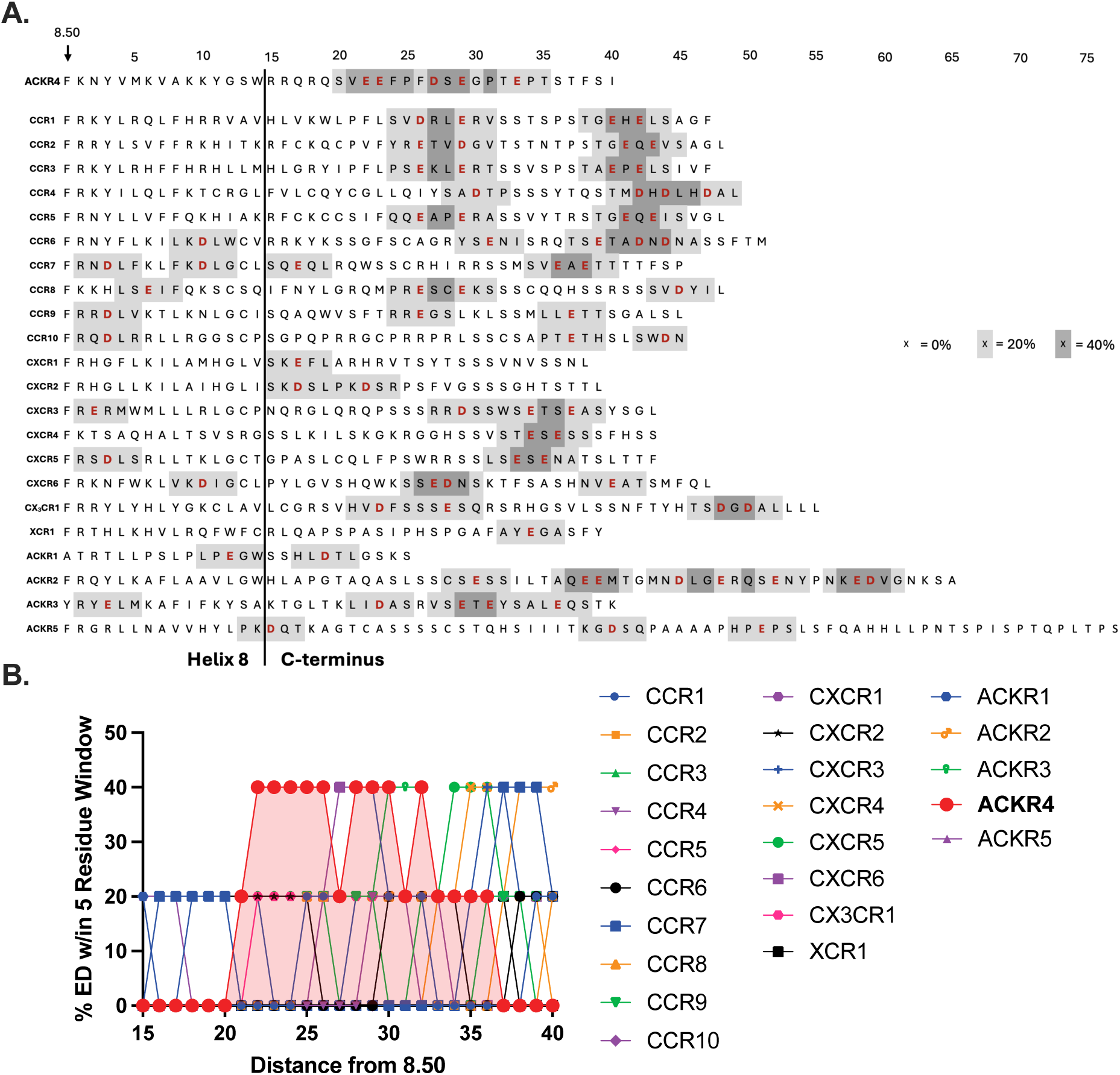
ACKR4 has a unique acidic-rich proximal C-terminus, compared to other chemokine receptors. (A) Sequence alignment of helix 8 and the C-terminus of all chemokine receptors. Percentage of acidic residues within the C-terminus of all chemokine receptors with a rolling analysis window of 5 residues are indicated, from the conserved aromatic residue at position 8.50. Sequence information was obtained from GPCRdb ^63^. (B) A sliding-window analysis (window size = 5 amino acids) was used to quantify the local density of acidic residues (Asp and Glu) in the C-termini of each chemokine receptor, reported in %. Higher values indicate short sequence segments that are enriched with negatively charged residues. ACKR4 is highlighted in red.

Acidic residues N-terminal to the target serines or threonines are necessary for GRK2/3 phosphorylation of peptide substrates ^46,47^. Therefore, we postulated that the uniquely dense acidic region preceding the majority of the ACKR4 phosphorylation sites may mediate the G protein-independent GRK2/3 activity. The identified glutamate and aspartate residues were mutated into alanines in pairs (E332A/E333A, D337A/E339A), in clusters (Proximal: E332A/E333A/D337A, Distal: E339A/E343A), or in their entirety (Acidic_All: E332A/E333A/D337A/E339A/E343A) (**Fig. 6A**). First, the effect of these mutations was assessed by measuring the arrestin recruitment by BRET. Constitutive arrestin engagement was reduced for all mutants to ∼60% of WT ACKR4 (**Fig. 6B**). Moreover, CCL25-mediated arrestin recruitment was nearly abolished when all five acidic sites were mutated (**Fig. 6C**). The Proximal sites had the next largest impact with ∼50% WT ACKR4 recruitment levels, while the Distal cluster substitution alone had no effect on arrestin engagement. No pair or cluster of residues was fully responsible for the impaired recruitment observed with the Acidic_All mutant. While these results are suggestive a role for the acidic motifs in mediating GRK2/3 recruitment and phosphorylation, an alternate explanation is that the residues impact the engagement of ACKR4 with the arrestins directly. To test that the mutational impact is on GRK2/3 recruitment, Gβγ was co-expressed to provide an alternate mechanism for membrane recruitment and bypass our proposed ACKR4 specific one. When Gβγ was included in the assay, arrestin recruitment of Acidic_All ACKR4 increased from <20% of WT to ∼50%, consistent with the interpretation that these residues are playing a role in coordinating GRK2/3 recruitment to the receptor rather than strictly impairing arrestin engagement (**Fig 6D**). The lack of full recovery could be partially explained by lower expression of the Acidic_All mutant relative to WT ACKR4 (**Supplemental Fig. 2**). We next tested the effect of the DE/A mutations on ACKR4-mediated GRK3 recruitment directly by bystander BRET between the kinase and mV-CAAX. Consistent with arrestin recruitment, GRK3 recruitment was severely reduced for the Acidic_All construct (**Fig. 6E and F**). The trend between the mutants was also similar to the arrestin recruitment, with the small exception that the Proximal substitution was as impaired as the Acidic_All rather than the next most impacted. While surface expression of all mutants was less than WT ACKR4, these differences did not correlate with GRK3 or arrestin recruitment and thus cannot explain the effects on effector recruitment observed here (**Supplemental Fig. 3**).

**Figure 6.**
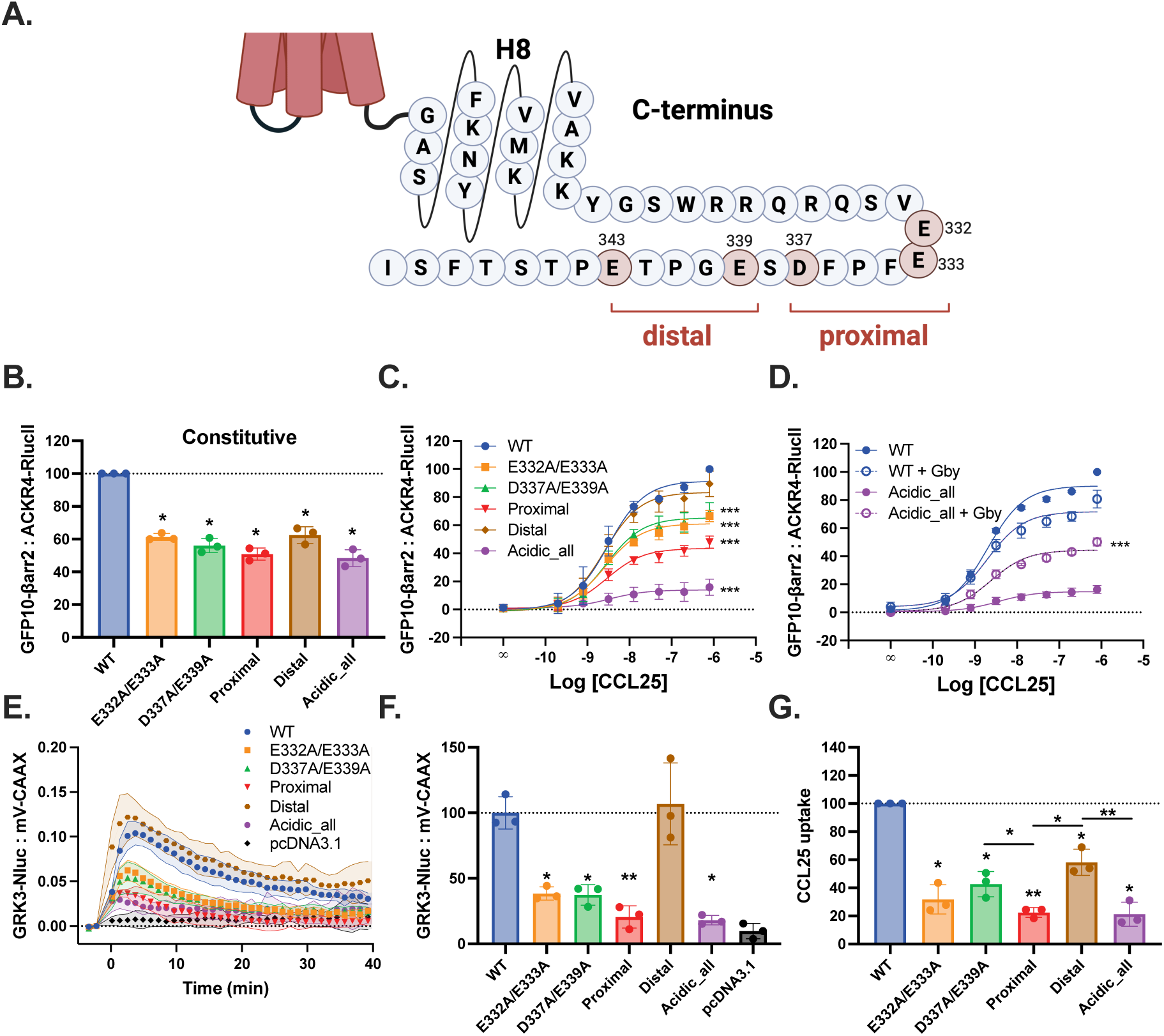
Acidic residues of ACKR4 facilitate GRK3 recruitment towards the membrane. (A) Schematic illustration of the C-terminus of ACKR4 with annotated acidic residues. Constitutive (B) and chemokine-induced (C) arrestin recruitment of GFP10-βarr2 towards ACKR4-RlucII, with co-expression of Gβγ (D), across a range of chemokine concentrations. (E) GRK3 localization measured as bystander BRET between GRK3-Nluc and mV-CAAX with co-expression of untagged ACKR4 over time after stimulation with 100 nM chemokine at time zero. (F) Quantified by area under the curve (AUC) from E. (G) Uptake of CCL25-AZ488 by ACKR4 measured by flow cytometry. Values represent the mean ± SD of three independent experiments performed in triplicate. Statistical significance at the top of the sigmoidal curve was determined by using the extra sum-of-squares F test, whereas significance for bar graphs was determined using Welch ANOVA followed by a Dunnett’s T3 multiple comparisons test (basal). ∗p < 0.05, ∗∗p < 0.001, and ∗∗∗p < 0.0001.

The role of ACKR4 is primarily to remove chemokines and sequester these ligands inside of cells ^8,9^. To quantify the effects of the acidic substitutions on CCL25 uptake, we measured the accumulation of fluorescent CCL25 into HEK293 cells and quantified the intensity of fluorescent emission after 1.5 h incubation by flow cytometry. Chemokine uptake by all of the DE/A mutants was significantly reduced compared to WT ACKR4 (**Fig. 6G**), matching the effects on GRK3 and arrestin recruitment for most of the substitutions. The decrease in uptake for the Distal construct matches the change in basal arrestin association, but not ligand induced, and suggests a role for receptor constitutive activity in chemokine scavenging. Like with GRK3 recruitment, the Proximal and Acidic_all substitutions had the greatest effects. Taken together, these results suggests that the acidic rich motif in the proximal ACKR4 C-terminus mediates GRK2/3 recruitment and chemokine scavenging functions.

### Basic residues in the GRK3 kinase domain coordinate recruitment to activated GPCRs

The noncanonical phosphorylation of ACKR4 by GRK3 appears to be coordinated by the acidic-rich C-terminus of the receptor (**Fig. 6**). This suggests that this unique motif may make distinct, ACKR4-specific interactions with the kinase. To explore this postulate, a model of the CCL25-ACKR4-GRK3 complex was generated with AlphaFold3 (**Fig. 7A**). In this model, the negatively-charged ACKR4 C-terminus is predicted near a series of basic residues along the kinase large lobe which may support coordinating the C-terminus in the kinase domain for phosphate modification. Since the predicted positioning of the C-terminus with respect to the kinase is of low confidence, specific interactions were not predicted (**Supplemental Fig. 4**). Seven lysine and arginine residues on GRK3 were mutated individually to alanines (R226A, K230A, R316A, K319A, K345A, K364A, and K383A) to eliminate potential electrostatic interactions with the acidic residues of the ACKR4 C-terminus and the recruitment of GRK3 to the membrane was measured by bystander BRET (**Fig. 7B**). While many of the mutations impaired GRK3 localization to ACKR4, these perturbations also similarly diminish GRK3 recruitment to the CCKRs CCR9 and CXCR4, suggesting that these putative interactions are not the unique, G protein-independent mechanism of ACKR4, but rather are generally involved for all GPCRs. Total expression of some of GRK mutants was slightly lower compared to WT, but do not explain the observed effects (**Supplemental Fig. 5**). This suggests the role of the basic residues on the GRK3 large lobe may play similar roles for coordinating GRK recruitment to ACKR4 as well as the G protein-dependent GPCRs. Based on the predicted positioning in the model, this would imply that the GPCR C-terminal interactions with GRK3 contribute to the recruitment of the kinase even under G protein-dependent conditions.

**Figure 7.**
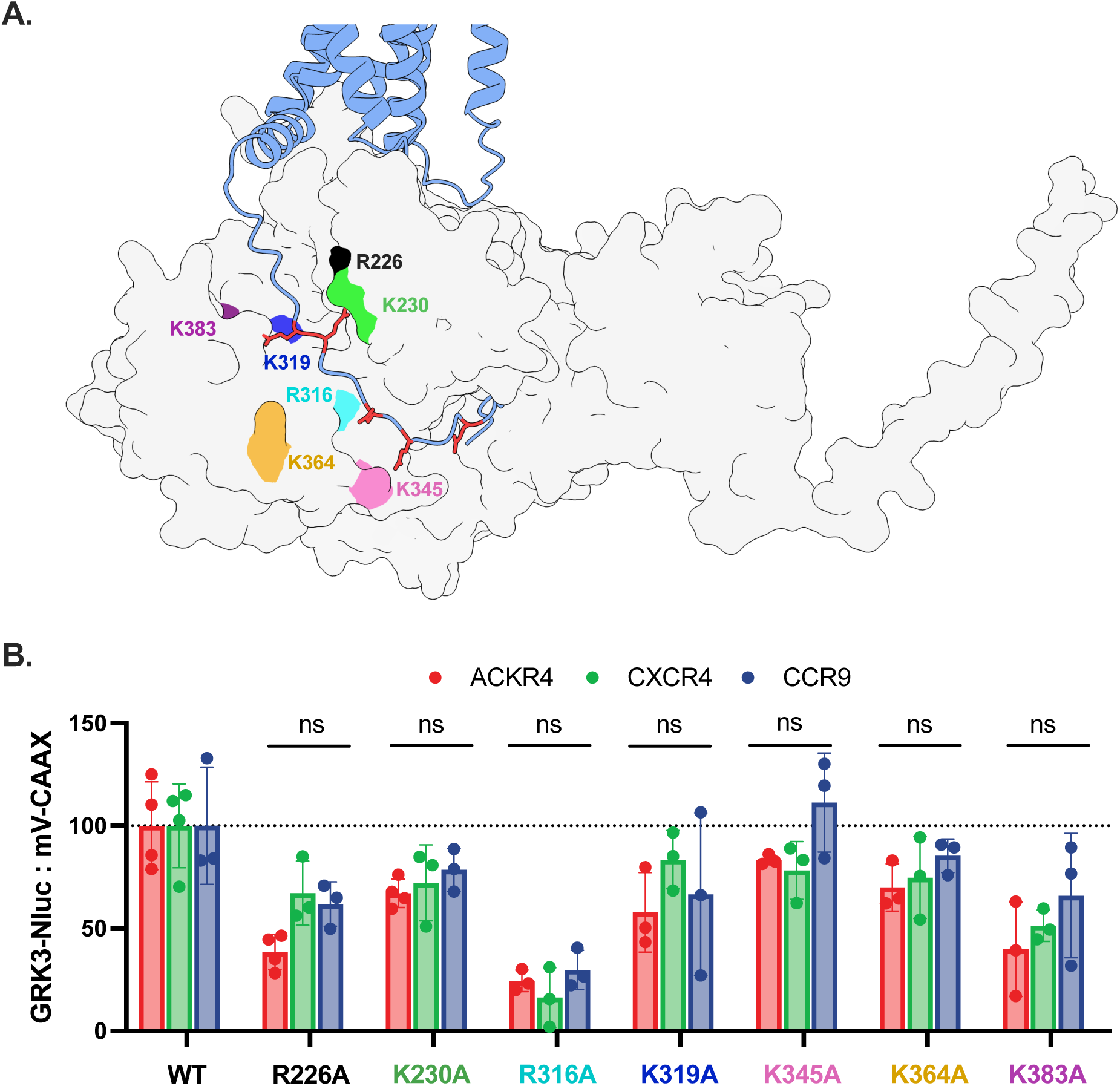
GRK3 binds similar to ACKR4 compared to CXCR4 and CCR9. (A) Alphafold3.0 model of ACKR4 with GRK3 with potential interacting basic residues highlighted, R226 -black, K230 -green, R316 -cyan, K319 -blue, K345 -pink, K364 -gold, and K383 -purple. The C-terminal acidic residues on ACKR4 are colored red. (B) GRK3 localization (GRK3-Nluc) towards the membrane (mV-CAAX) quantified by AUC. Values represent the mean ± SD of three independent experiments performed in triplicate. Statistical significance was determined by one-way Brown–Forsythe and Welch ANOVA followed by a Dunnett’s T3 multiple comparisons test.

The model (**Fig. 7A**) suggests that the acidic residues may play a role in orienting the C-terminus for phosphate addition. Thus, another key component to this interaction would be the catalytic site of the kinase with the to-be-modified serine or threonine residue. Contacts during the enzymatic reaction may contribute to the GRK recruitment to ACKR4. While these interactions would be present for all GPCRs, they may be relatively minor compared to canonical G protein-mediated translocation of GRK2/3 and would represent a larger source of binding energy for the ACKR4 mechanism. To interrogate the role of these interactions on GRK recruitment, a point mutation was added to GRK3 within the ATP-binding pocket, K220R, which renders the kinase ‘kinase-dead’ and unable to phosphorylate substrates ^48^. Recruitment of the kinase-dead GRK3 (KD-GRK3) was tracked by membrane translocation by bystander BRET between GRK3-Nluc and mV-CAAX. Following stimulation with CCL25, the recruitment of KD-GRK3 to ACKR4 is only about 25% compared to the WT GRK3 (**Fig. 8A**). None of the receptors which depend on G proteins for GRK3 recruitment, ACKR3, CXCR4, or CCR9, showed significant differences between WT and KD kinase recruitment (**Fig. 8B, C, and D**). Total expression of the KD-GRK3-Nluc, measured by luminescence counts, was similar compared to WT GRK3 (**Supplemental Fig. 6**). The contribution of functional phosphorylation on ACKR4 is further supported by ∼40% GRK3 recruitment to an ACKR4 construct with all potentially phosphorylated serine and threonine residues substituted by alanine (**Supplemental Fig. 7A and B**). Surface expression of the ST/A mutant was slightly less than WT ACKR4 (**Supplemental Fig. 7C**). This suggests that, when GRK2/3 recruitment is independent of G protein interactions, kinase recruitment is governed by the interactions responsible for positioning and facilitating the phosphorylation reaction.

**Figure 8.**
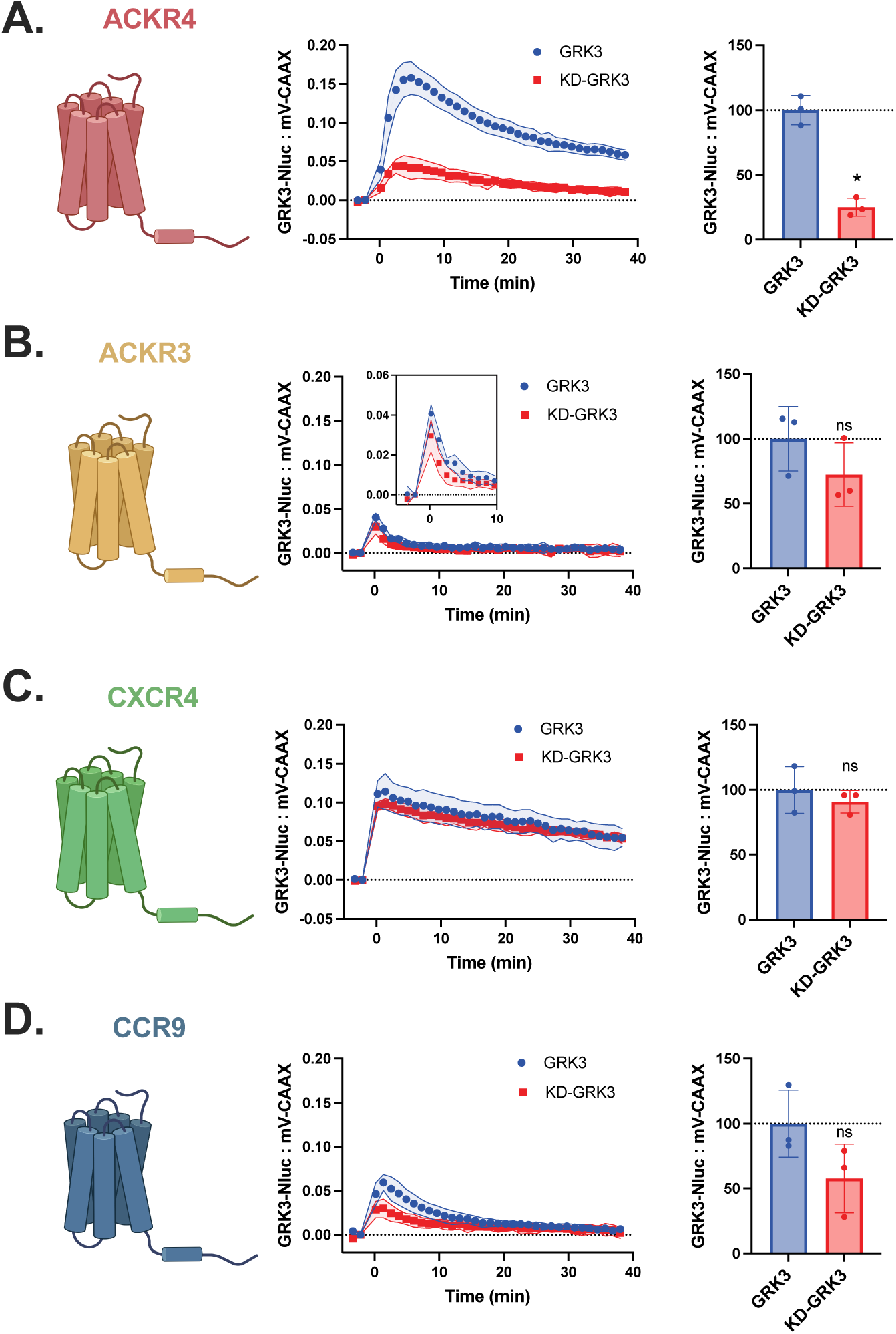
Functional kinase is required for ACKR4 recruitment of GRK3. GRK3 localization towards the membrane measured as bystander BRET between GRK3-Nluc and mV-CAAX with co-expression of untagged ACKR4 (A), ACKR3 (B), CXCR4 (C), or CCR9 (D) following stimulation with 100 nM CCL25 (ACKR4 and CCR9) or CXCL12 (ACKR3 and CXCR4). Quantification (right) by AUC. Values represent the mean ± SD of three independent experiments performed in triplicate. Statistical significance was determined by one-way Brown–Forsythe and Welch ANOVA followed by a Dunnett’s T3 multiple comparisons test (basal). ∗p < 0.001.

## Discussion

ACKR4 plays a crucial role in modulating chemokine gradients by scavenging the chemokines CCL19, CCL20, CCL21, and CCL25 to regulate the migration of CCR6-, CCR7-, and CCR9-expressing immune cells. This activity is mediated by arrestin-coupling and GRK phosphorylation. Specifically, GRK2/3 dominate ACKR4 phospho-modification, creating a paradox. GRK2/3 require interactions with activated G proteins to translocate to the PM and phosphorylate GPCRs, however, ACKR4 does not activate G proteins. In fact, ACKR4 does not need G proteins to recruit the G protein-dependent kinases, but rather coordinates the enzymes through a uniquely acidic-rich proximal C-terminus. Interactions with these residues, along with specific sites for phosphorylation, provide sufficient binding energy to bypass the canonical G protein interactions. These findings highlight a unique mechanism by which an atypical GPCR has evolved to utilize an efficient regulation system despite its atypical function (**Fig. 9**).

**Figure 9.**
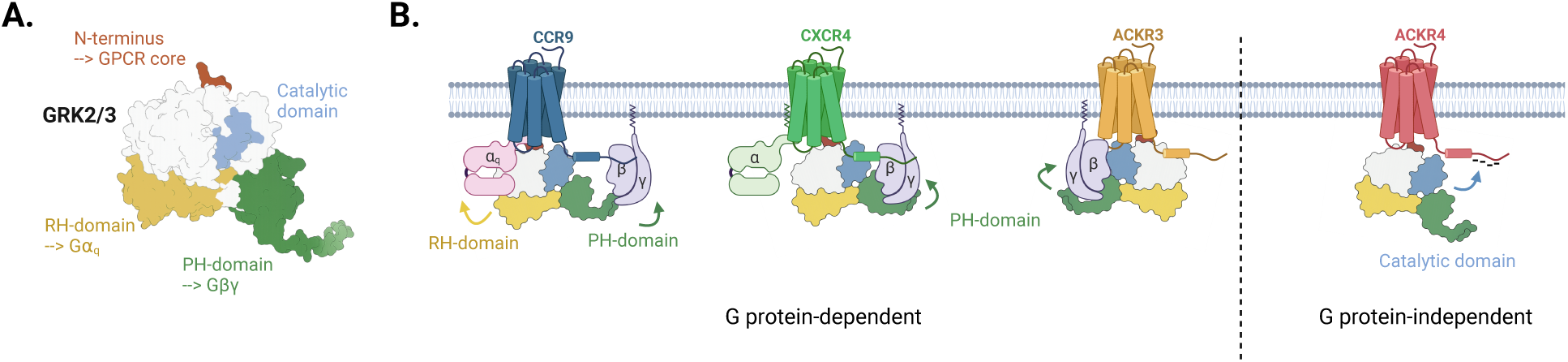
The ACKR4 coordinates GRK2/3 recruitment by interactions between the acidic-rich C-terminus and kinase catalytic domain. (A) Phosphorylation of GPCRs by GRK2/3 is coordinated by protein-protein interactions which promote PM localization by complexing with the activated Gβγ (PH-domain, Green) and/or Ga_q_ (RH-domain, Yellow) subunits. Active receptors are identified by the kinases through the GRK N-terminus (Orange). The GRK catalytic domain (Blue) then binds the substrate C-terminus and modifies specific serine and/or threonine residues. (B) Most GPCRs require the G protein interactions to relocate the GRKs to the PM for the GPCR specific interactions and phosphorylation. ACKR4 can utilize GRK2/3 without G protein coordination and only requires interactions with the receptor core (GRK N-terminus) and C-terminal tail (GRK catalytic domain).

The preference of GRK2/3 to phosphorylate serine and threonine residues C-terminal following acidic motifs is well documented ^46^. For example, mutation of a glutamate to a lysine before the terminal phosphosites in CXCR4 impairs phosphorylation of these positions and protects the receptor from desensitization ^49^. Similarly, GRK2 readily phosphorylates peptides derived from phosphorylation motifs within the ⍺2-adrenergic receptor, but only when the target serine or threonine is preceded by an acidic residue ^47^. A single glutamate is sufficient for GRK2 activity, with charges two or three residues before the phosphosite being the most efficient sequences. Many of the putative phosphorylation sites on ACKR4 are within three residues of an N-terminal acidic residue, suggesting the receptor may have evolved these acidic motifs to allow the C-terminus to act as an efficient substrate for GRK2/3 to remove the G protein dependency.

Despite efficient coupling of the kinase domain to the substrate’s C-terminus, GRK2/3 phosphorylation of ACKR4 is ultimately still dependent on receptor activation. This observation is consistent with earlier reports showing a 200-fold increase in efficiency of free peptide phosphorylation by GRK2 when an activated GPCR, either light-activated rhodopsin or agonist-activated β_2_AR, was included ^50^. Thus, even in the absence of G protein dependency, receptor activation coordinates phosphorylation with chemokine detection and subsequent scavenging. Although the explicit role of phosphorylation for chemokine uptake has not yet been reported, arrestins play a clear role in efficient chemokine clearance for ACKR4 ^17^. Therefore, the detection of the active ACKR4 state by GRK2/3 allows for synchronization of chemokine binding with arrestin coupling despite lacking the second level of regulation through G proteins.

While seven GRKs are expressed in the human genome, the vast majority of GPCR regulation is performed by the four ubiquitously expressed kinases, GRK2/3/5/6. The levels of these kinases can vary drastically by cell and provide cell-type specific, tailored signaling responses from different GPCRs. ACKR4 is expressed on a variety of cell types including lymphatic endothelial cells ^51^, thymic epithelial cells ^52^, and subpopulations of gut-derived mesenchymal stem cells ^53^. In each environment, the atypical receptor functions to regulate the surrounding chemokine gradients and immune cell positioning and responses. Phosphorylation and arrestin coupling contribute to chemokine scavenging by ACKR4 ^17^, thus, the evolved GRK2/3 phosphorylation mechanism would allow the receptor to utilize all ubiquitously-expressed GRKs and efficiently regulate chemokine availability across an array of cellular backgrounds.

Five ACKRs have been identified that have evolved to function independently of G proteins. ACKR1 functions independent of canonical effects such as G protein or arrestins and shows no discernable recruitment of any effectors, including GRKs, upon chemokine stimulation ^54^. The distal C-terminus of ACKR2 is similarly rich with acidic residues interspersed with putative phosphorylation sites. The positioning of these motifs could facilitate similar G protein-independent GRK2/3 phosphorylation as described here for ACKR4. Deletion of these residues eliminates agonist-induced arrestin recruitment and internalization ^55^. ACKR3 phosphorylation is primarily dependent on GRK5/6, but can borrow Gβγ from other sources, such as co-activation of CXCR4, to drive GRK2/3-mediated phosphorylation ^26^. ACKR5 is solely dependent on GRK5/6 for its ligand-induced regulation and knocking out GRK2/3 had no effect on ligand-induced receptor internalization and uptake ^56^. As described here, ACKR4 has evolved a C-terminus to be an efficient enough substrate for GRK2/3 phosphorylation to overcome G protein-dependency for membrane targeting. These distinct mechanisms allow for ACKRs to circumvent the need for G protein activation while still utilizing the evolved GRK regulatory mechanisms.

Taking the putative phosphorylation sites alone, ACKR4 lacks a known arrestin-binding motif, despite being an arrestin-biased, atypical receptor. Neither the simplified **PXPP** motif ^57^ nor the extended **PX_1-2_PXXP** motif ^58^, where **P** is a phosphorylated serine or threonine and **X** is any amino acid, are present in the sequence. Only when the glutamates and aspartates of the acidic-rich proximal C-terminus are treated as phosphomimetics are clear arrestin-binding sequences apparent. By replacing one or more of the **P**’s from the binding sequence with a negatively charged residue, it would effectively render part of the binding barcode always ‘on’ and reduce the needed modifications for arrestin recruitment. These primed sequences could respond quicker to chemokine stimulation, perhaps allowing for rapid changes in chemokine concentration to be efficiently rectified.

Treating the acidic motifs as phosphomimetics also suggests that phosphorylation along the C-terminus could similarly promote further phosphorylation of downstream sites. Peptide phosphorylation experiments indicate that phosphorylated serines N-terminal to the target residue enhance the rate and efficiency of further GRK2 phosphorylation ^50^. Many examples of hierarchal phosphorylation have also been reported. CXCR4 ^49^, C5a anaphylatoxin receptor ^59^, rhodopsin ^60^, and β_1_/β_2_AR ^61^ all show sequential and ordered phosphate incorporation. Phosphorylation of the µ-opioid receptor at S375 is required before either T376 or T379 can be modified ^62^. This subsequent phosphorylation is selectively mediated by GRK2/3, consistent with the kinase preference for N-terminal negative charges. Our results here suggest that the G protein-dependence of GRK2/3 could become less as more phosphates are transferred to the receptors, which may allow for rapid saturation of the GPCR C-terminus and efficient signal termination.

In conclusion, ACKR4 non-canonically recruits the GRK2/3 family that is driven by its acidic proximal C-terminus. The acidic residues coordinate the kinase domain to orient the target serine or threonine residues for phosphorylation and these interactions are sufficient to overcome the lack of G protein coupling and coordination. This unique feature allows the atypical receptor to exploit the efficient phosphorylation by GRK2/3 without triggering of G protein signaling cascades. Such a mechanism may be more broadly applicable to other GPCRs as a secondary regulatory motif after G proteins and provides important insights into ACKR4 regulation.

## Supporting information

This article contains supporting information.

## Acknowledgements

We thank C. Hoffmann (Friedrich-Schiller-Universität) for the BRET constructs and GRK K/O cell lines used in this study.

## Data availability

The authors declare that all the data supporting the findings of this study are available within the paper and its Supplemental Data.

## Author contribution

T.D.L. and C.T.S. conceptualization; T.D.L. and C.T.S. formal analysis; M.J.S and C.T.S. funding acquisition; T.D.L. and I.B.S.A. investigation; T.D.L and C.T.S. methodology; C.T.S. project administration; M.J.S. and C.T.S. resources; C.T.S. supervision; T.D.L and C.T.S. validation; T.D.L and C.T.S. visualization; T.D.L and C.T.S. writing – original draft; T.D.L., I.B.S.A., M.J.S., and C.T.S. writing – review & editing.

## Funding and additional information

This publication is part of the TRANSLATION project with file number OCENW.M.24.006 which is partly financed by the Dutch Research Council (NWO) under the grant [grant ID: https://doi.org/10.61686/YKUZU08217].

## Nonstandard Abbreviations

ACKR: atypical chemokine receptor
ACKR3: atypical chemokine receptor 3
ACKR4: atypical chemokine receptor 4
AUC: area under curve
BRET: bioluminescence resonance energy transfer
CCL25: C-C chemokine 25
CXCL12: C-X-C chemokine 12
CCKR: canonical chemokine receptor
CCR9: C-C chemokine receptor 9
CXCR4: C-X-C chemokine receptor 4
GPCR: G protein-coupled receptor
GRK: GPCR kinase
WT: wild type
rGFP: *Renilla* green fluorescent protein
RlucII: *Renilla* luciferase II
PM: plasma membrane
mV: monomeric venus fluorescent protein
KD: kinase dead

