## Supplemental Information for "GPCR kinase 3 phosphorylates atypical chemokine receptor 4 independent of G proteins"

### Supplementary Information

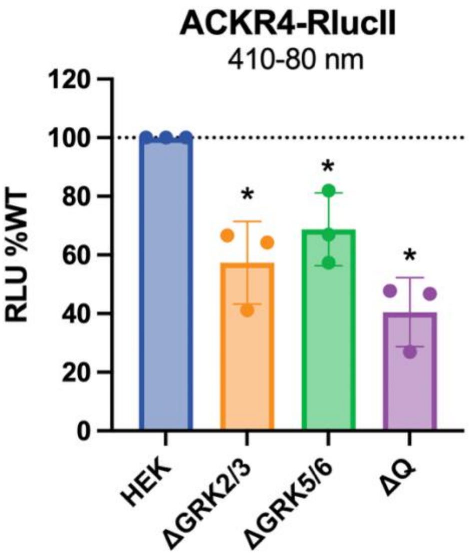


**Supplementary Figure 1. ACKR4 expression in HEK293 parental cells and GRK knockouts.** Total expression of ACKR4-RlucII measured by luminescence counts (410-80 nm) in HEK293, ΔGRK2/3, ΔGRK5/6, and ΔQ cells. Values represent the mean ± SD of three independent experiments performed in triplicate normalized to WT luminescence. Statistical significance compared to parental cells was determined by one-way Brown–Forsythe and Welch ANOVA followed by a Dunnett's T3 multiple comparisons test. *$p < 0.05$

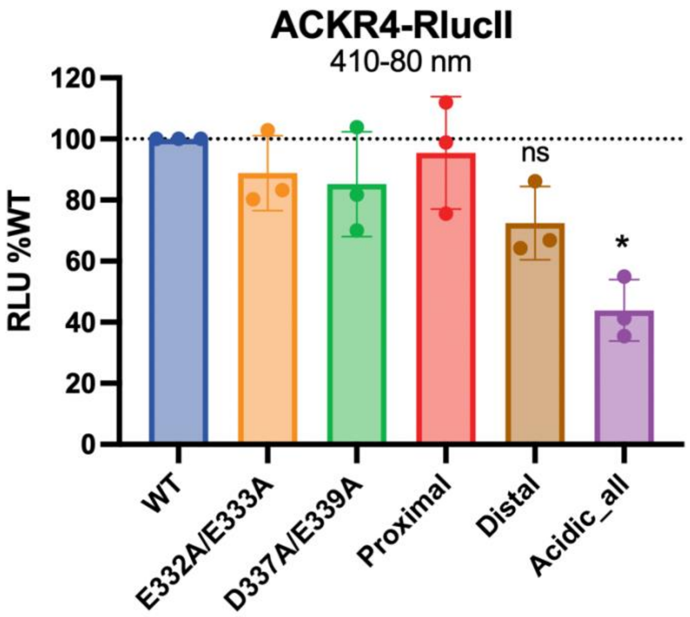


**Supplementary Figure 2. ACKR4 expression levels of DE/A mutants in HEK293 cells.** Total expression of ACKR4-RlucII measured by luminescence counts (410-80 nm). Values represent the mean ± SD of three independent experiments performed in triplicate normalized to WT luminescence. Statistical significance compared to WT ACKR4 expression was determined by one-way Brown–Forsythe and Welch ANOVA followed by a Dunnett's T3 multiple comparisons test. *$p < 0.05$

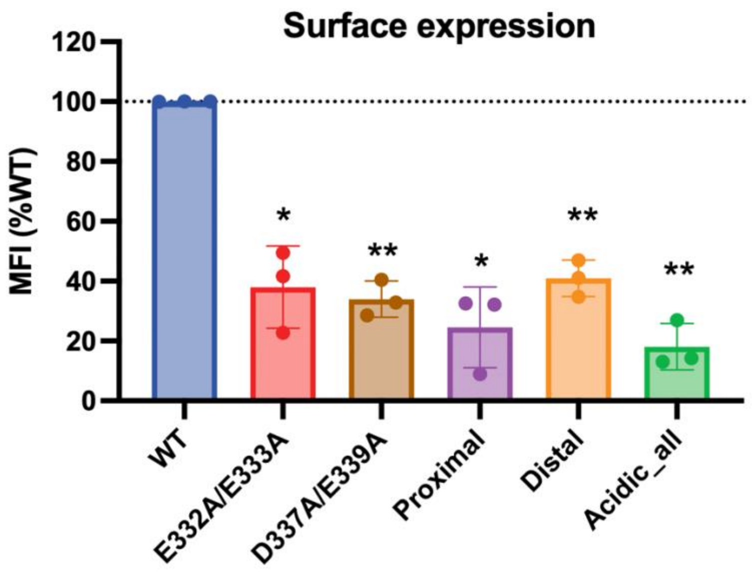


**Supplementary Figure 3. ACKR4 surface expression levels of DE/A mutants in HEK293 cells.** Surface expression of untagged ACKR4 measured by flow cytometry. Values represent the mean ± SD of three independent experiments performed in triplicate normalized to WT mean fluorescence intensity (MFI). Statistical significance was determined by one-way Brown–Forsythe and Welch ANOVA followed by a Dunnett's T3 multiple comparisons test. $*p < 0.05$, $**p<0.001$

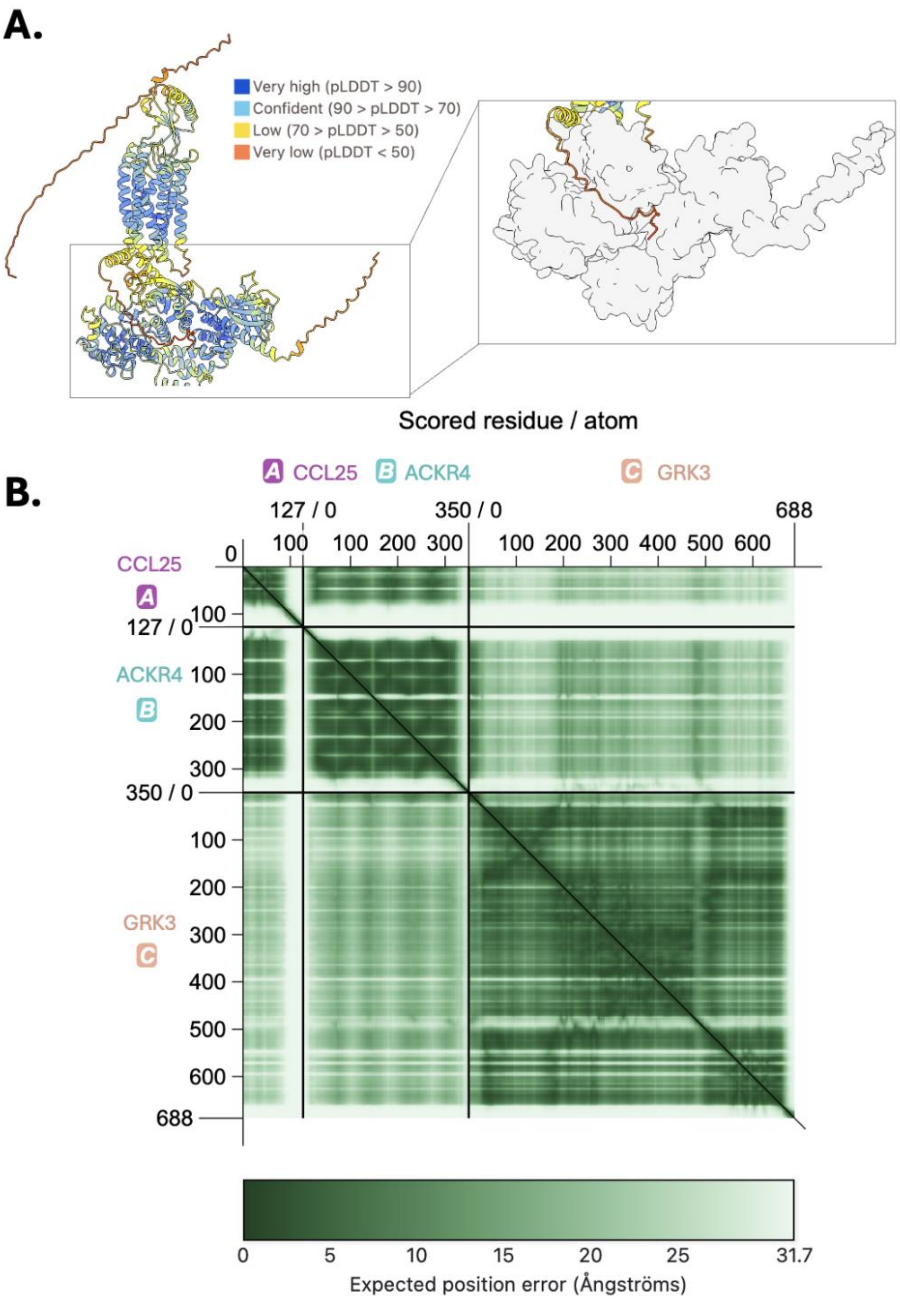


**Supplementary Figure 4. Predicted ACKR4 C-terminal orientation near the kinase domain.** (A) Alphafold3.0 model of the CCL25-ACKR4-GRK3 complex with predicted local distance difference test (pLDDT) per residue. (Inset) C-terminus of ACKR4 inside the pocket of GRK3. (B) PAE plot of CCL25 [A], ACKR4 [B], GRK3 [C]. Plots are generated by PAE Viewer [1].

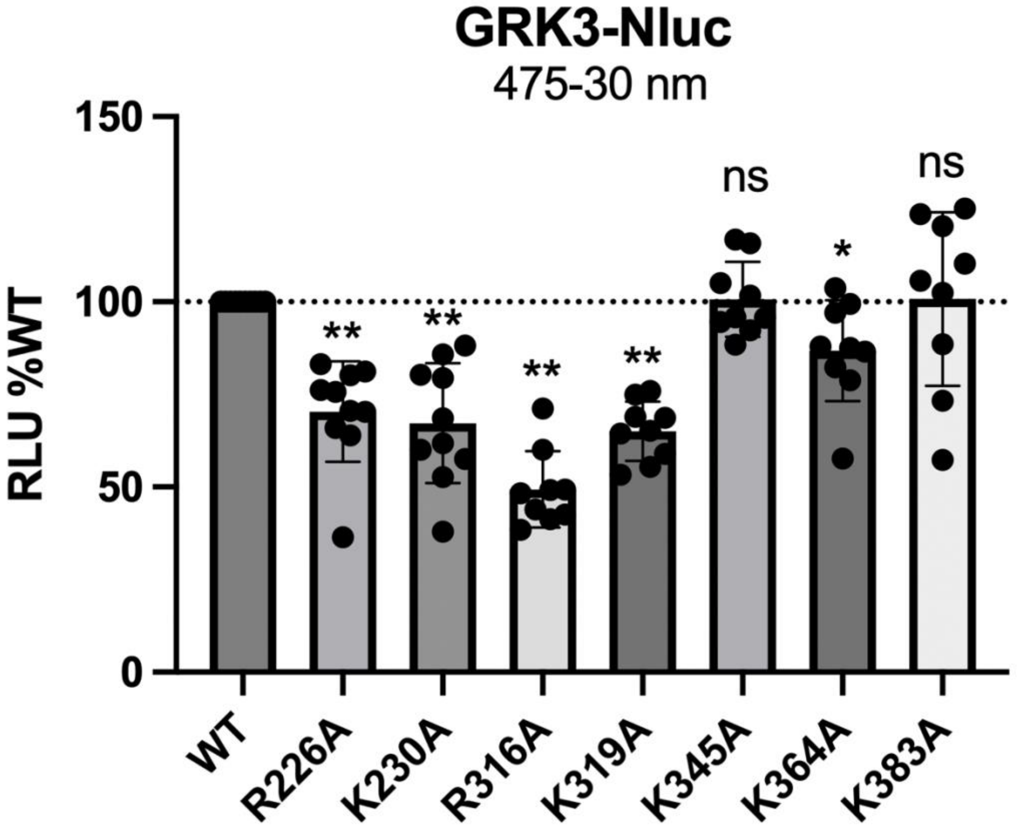


**Supplementary Figure 5. GRK3 expression levels in HEK293 cells.** Total expression of GRK3-Nluc measured by luminescence counts (475-30 nm). Values represent the mean ± SD of three independent experiments (per GPCR) performed in triplicate normalized to WT luminescence. Statistical significance comparing the expression of mutant GRKs to WT GRK3 was determined by one-way Brown–Forsythe and Welch ANOVA followed by a Dunnett's T3 multiple comparisons test. *$p < 0.001$, and **$p < 0.0001$.

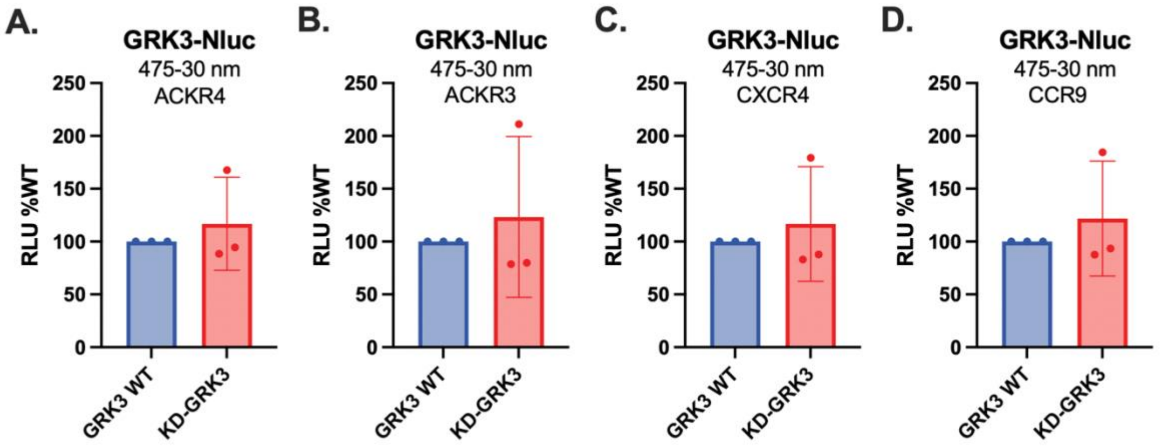


**Supplementary Figure 6. (KD-)GRK3 expression levels in HEK293 cells.** Total expression of GRK3-Nluc measured by luminescence counts (475-30 nm). Bars represent the mean ± SD of three independent experiments (points) performed in triplicate normalized to WT luminescence.

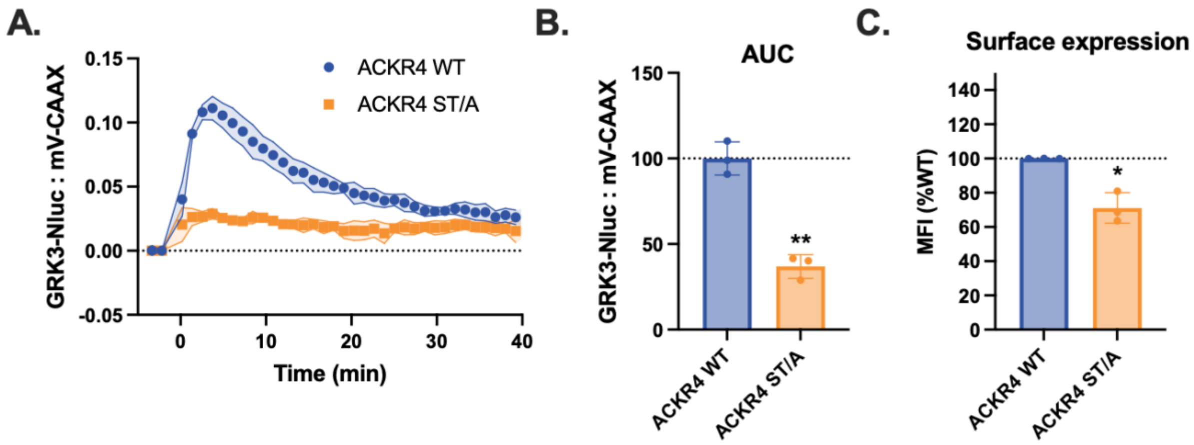


**Supplementary Figure 7. ST/A mutation impairs ACKR4-mediated GRK3 localization towards the membrane.** (A) GRK3 localization towards the membrane measured as bystander BRET between GRK3-Nluc and mV-CAAX with co-expression of untagged WT ACKR4 and ACKR4 ST/A (S330A/S338A/T342A/T345A/S346A/T347A/S349A). Transfected DNA for ACKR4 ST/A was increased to match surface expression of WT. (B) Quantification by AUC. (C) Surface expression of untagged ACKR4 measured by flow cytometry. Values represent the mean ± SD of three independent experiments performed in triplicate. Statistical significance was determined by one-way Brown–Forsythe and Welch ANOVA followed by a Dunnett's T3 multiple comparisons test (basal). *$p < 0.05$, **$p < 0.001$.

### Supplementary Table 1. Statistics

| Fig. | Condition | Emax | | Bar graph | | Statistical test |
|---|---|---|---|---|---|---|
| | | P-value | Value ± SD/CI* | P-value | Value ± SD | |
| 1c | HEK P | N.A | 86.33 ± 5.08* | N.A. | | Extra sum-of-squares F test |
| | ΔGRK2/3 | <0.0001 | 28.13 ± 2.89* | | | |
| | ΔGRK5/6 | <0.0001 | 62.94 ± 4.91* | | | |
| | ΔQ | <0.0001 | 3.37 ± 0.54* | | | |
| 2b-L | - | N.A. | 0.122 ± 0.014 | N.A. | | t-test |
| | +GRK3-CT | 0.43 | 0.145 ± 0.038 | | | |
| 2c-L | - | N.A. | 0.044 ± 0.0098 | N.A. | | t-test |
| | +GRK3-CT | 0.019 | 0.013 ± 0.0039 | | | |
| 2d-L | - | N.A. | 0.12 ± 0.013 | N.A. | | t-test |
| | +GRK3-CT | 0.0096 | 0.056 ± 0.0038 | | | |
| 2e-L | - | N.A. | 0.059 ± 0.010 | N.A. | | t-test |
| | +GRK3-CT | 0.017 | 0.0090 ± 0.016 | | | |
| 2b-R | - | N.A. | N.A. | N.A. | | Extra sum-of-squares F test |
| | +GRK3-CT | N.A. | N.A. | | | |
| | +Gby | N.A. | N.A. | | | |
| 2c-R | - | N.A. | 98.02 ± 4.58 | N.A. | | Extra sum-of-squares F test |
| | +GRK3-CT | <0.0001 | 63.69 ± 3.43 | | | |
| | +Gby | <0.0001 | 123.7 ± 1.80 | | | |
| 2d-R | - | N.A. | 93.49 ± 4.30 | N.A. | | Extra sum-of-squares F test |
| | +GRK3-CT | <0.0001 | 65.71 ± 4.01 | | | |
| | +Gby | N.A. | N.A. | | | |
| 2e-R | - | N.A. | 101.2 ± 2.21 | N.A. | | Extra sum-of-squares F test |
| | +GRK3-CT | <0.0001 | 89.03 ± 2.97 | | | |

| | | | | | | |
|---|---|---|---|---|---|---|
| | +Gby | N.A. | N.A. | | | |
| **3b-L** | ΔQ | N.A. | | N.A. | 0.49 ± 0.05 | Browns-Forsythe & Welch ANOVA |
| | ΔQ+GRK2 | | | 0.004 | 2.05 ± 0.09 | |
| | ΔQ+GRK2 R587Q | | | 0.017 | 0.90 ± 0.09 | |
| | ΔQ+GRK3 | | | 0.004 | 2.11 ± 0.10 | |
| | ΔQ+GRK3 R587Q | | | 0.0023 | 0.90 ± 0.06 | |
| **3c-L** | ΔQ | N.A. | | N.A. | 0.91 ± 0.04 | Browns-Forsythe & Welch ANOVA |
| | ΔQ+GRK2 | | | 0.040 | 1.67 ± 0.17 | |
| | ΔQ+GRK2 R587Q | | | 0.96 | 0.92 ± 0.04 | |
| | ΔQ+GRK3 | | | 0.002 | 1.78 ± 0.09 | |
| | ΔQ+GRK3 R587Q | | | 0.95 | 0.92 ± 0.05 | |
| **3d-L** | ΔQ | N.A. | | N.A. | N.A. | Browns-Forsythe & Welch ANOVA |
| | ΔQ+GRK2 | | | N.A. | N.A. | |
| | ΔQ+GRK2 R587Q | | | N.A. | N.A. | |
| | ΔQ+GRK3 | | | N.A. | N.A. | |
| | ΔQ+GRK3 R587Q | | | N.A. | N.A. | |
| **3f-L** | ΔQ | N.A. | | N.A. | 0.47 ± 0.02 | Browns-Forsythe & Welch ANOVA |
| | ΔQ+GRK2 | | | 0.0007 | 1.95 ± 0.09 | |
| | ΔQ+GRK2 R587Q/D110A | | | 0.003 | 0.90 ± 0.04 | |
| | ΔQ+GRK3 | | | 0.0001 | 1.94 ± 0.05 | |
| | ΔQ+GRK3 | | | 0.018 | 0.74 ± 0.08 | |

| | | | | | | |
|---|---|---|---|---|---|---|
| | R587Q/D110A | | | | | |
| **3g-L** | ΔQ | N.A. | | N.A. | N.A. | Browns-Forsythe & Welch ANOVA |
| | ΔQ+GRK2 | | | N.A. | N.A. | |
| | ΔQ+GRK2 R587Q/D110A | | | N.A. | N.A. | |
| | ΔQ+GRK3 | | | N.A. | N.A. | |
| | ΔQ+GRK3 R587Q/D110A | | | N.A. | N.A. | |
| **3b-R** | ΔQ | N.A. | N.A. | N.A. | | Extra sum-of-squares F test |
| | ΔQ+GRK2 | N.A. | 62.05 ± 4.73 | | | |
| | ΔQ+GRK2 R587Q | <0.0001 | 90.48 ± 5.55 | | | |
| | ΔQ+GRK3 | N.A. | 60.79 ± 5.90 | | | |
| | ΔQ+GRK3 R587Q | <0.0001 | 102.4 ± 5.11 | | | |
| **3c-R** | ΔQ | N.A. | N.A. | N.A. | | Extra sum-of-squares F test |
| | ΔQ+GRK2 | N.A. | 88.59 ± 4.02 | | | |
| | ΔQ+GRK2 R587Q | <0.0001 | 38.41 ± 2.97 | | | |
| | ΔQ+GRK3 | N.A. | 83.51 ± 3.72 | | | |
| | ΔQ+GRK3 R587Q | <0.0001 | 43.33 ± 2.01 | | | |
| **3d-R** | ΔQ | N.A. | N.A. | N.A. | | Extra sum-of-squares F test |
| | ΔQ+GRK2 | N.A. | 77.57 ± 5.87 | | | |
| | ΔQ+GRK2 R587Q | <0.0001 | 25.90 ± 2.95 | | | |
| | ΔQ+GRK3 | N.A. | 78.57 ± 5.12 | | | |
| | ΔQ+GRK3 | <0.0001 | 41.55 ± 2.53 | | | |

| | | | | | | |
|---|---|---|---|---|---|---|
| | R587Q | | | | | |
| **3f-R** | ΔQ | N.A. | N.A. | N.A. | | Extra sum-of-squares F test |
| | ΔQ+GRK2 | N.A. | N.A. | | | |
| | ΔQ+GRK2 R587Q/D110A | N.A. | N.A. | | | |
| | ΔQ+GRK3 | N.A. | N.A. | | | |
| | ΔQ+GRK3 R587Q/D110A | N.A. | N.A. | | | |
| **3g-R** | ΔQ | N.A. | N.A. | N.A. | | Extra sum-of-squares F test |
| | ΔQ+GRK2 | N.A. | 86.65 ± 4.34 | | | |
| | ΔQ+GRK2 R587Q/D110A | <0.0001 | 51.73 ± 2.56 | | | |
| | ΔQ+GRK3 | N.A. | 88.72 ± 4.50 | | | |
| | ΔQ+GRK3 R587Q/D110A | <0.0001 | 50.85 ± 3.65 | | | |
| **4b** | ΔQ | N.A. | | N.A. | 0.86 ± 0.04 | Browns-Forsythe & Welch ANOVA |
| | ΔQ+GRK2 | | | <0.0001 | 1.72 ± 0.05 | |
| | ΔQ+GRK2 R587Q | | | 0.22 | 0.95 ± 0.05 | |
| | ΔQ+GRK3 | | | 0.001 | 1.95 ± 0.01 | |
| | ΔQ+GRK3 R587Q | | | 0.16 | 0.95 ± 0.04 | |
| **4c** | ΔQ | N.A. | N.A. | N.A. | | Extra sum-of-squares F test |
| | ΔQ+GRK2 | N.A. | N.A. | | | |
| | ΔQ+GRK2 R587Q | N.A. | N.A. | | | |
| | ΔQ+GRK3 | N.A. | N.A. | | | |
| | ΔQ+GRK3 | N.A. | N.A. | | | |

| | | | | | | |
|---|---|---|---|---|---|---|
| | R587Q | | | | | |
| **6a** | WT | N.A. | | N.A. | N.A. | Browns-Forsythe & Welch ANOVA |
| | E332A/E333A | | | 0.001 | 61.31 ± 2.2 | |
| | D337A/E339A | | | 0.003 | 56.10 ± 4.4 | |
| | Proximal | | | 0.002 | 50.87 ± 3.7 | |
| | Distal | | | 0.006 | 62.47 ± 5.1 | |
| | Acidic_all | | | 0.003 | 48.43 ± 5.1 | |
| **6b** | WT | N.A. | 91.55 ± 5.51 | N.A. | | Extra sum-of-squares F test |
| | E332A/E333A | <0.0001 | 61.22 ± 3.37 | | | |
| | D337A/E339A | <0.0001 | 65.37 ± 3.84 | | | |
| | Proximal | <0.0001 | 43.55 ± 3.01 | | | |
| | Distal | N.A. | N.A. | | | |
| | Acidic_all | 0.0045 | 13.93 ± 2.91 | | | |
| **6c** | WT | N.A. | 90.11 ± 3.62 | N.A. | | Extra sum-of-squares F test |
| | WT + Gβγ | <0.0001 | 71.72 ± 4.14 | | | |
| | Acidic_all | N.A. | 14.82 ± 1.62 | | | |
| | Acidic_all + Gβγ | <0.0001 | 44.41 ± 2,61 | | | |
| **6e** | WT | N.A. | | N.A. | 100 ± 12.3 | Browns-Forsythe & Welch ANOVA |
| | E332A/E333A | | | 0.015 | 38.5 ± 5.08 | |
| | D337A/E339A | | | 0.019 | 37.4 ± 7.84 | |
| | Proximal | | | 0.003 | 20.6 ± 8.55 | |
| | Distal | | | 1.00 | 106.8 ± 31.3 | |
| | Acidic_all | | | 0.024 | 18.1 ± 3.55 | |
| **6f** | WT | N.A. | | N.A. | N.A. | Browns-Forsythe & Welch ANOVA |
| | E332A/E333A | | | 0.02 | 31.8 ± 10.4 | |
| | D337A/E339A | | | 0.02 | 42.7 ± 9.0 | |
| | Proximal | | | 0.002 | 22.4 ± 3.5 | |

| | | | | | | |
|---|---|---|---|---|---|---|
| | Distal | | | 0.04 | 58.2 ± 9.3 | |
| | Acidic_all | | | 0.01 | 21.3 ± 8.6 | |
| **7b** | WT | ACKR4 | N.A. | N.A. | N.A. | Browns-Forsythe & Welch ANOVA |
| | | CXCR4 | | | N.A. | |
| | | CCR9 | | | N.A. | |
| | R226A | ACKR4 | N.A. | 0.057 | 38.6 ± 8.5 | Browns-Forsythe & Welch ANOVA |
| | | CXCR4 | | | 67.2 ± 15.7 | |
| | | CCR9 | | | 61.8 ± 10.8 | |
| | K230A | ACKR4 | N.A. | 0.057 | 67.1 ± 7.0 | Browns-Forsythe & Welch ANOVA |
| | | CXCR4 | | | 72.2 ± 18.6 | |
| | | CCR9 | | | 78.5 ± 10.4 | |
| | R316A | ACKR4 | N.A. | 0.37 | 25.5 ± 5.3 | Browns-Forsythe & Welch ANOVA |
| | | CXCR4 | | | 16.3 ± 14.5 | |
| | | CCR9 | | | 29.8 ± 9.5 | |
| | K319A | ACKR4 | N.A. | 0.55 | 57.8 ± 19.3 | Browns-Forsythe & Welch ANOVA |
| | | CXCR4 | | | 83.5 ± 14.2 | |
| | | CCR9 | | | 66.6 ± 39.7 | |
| | K345A | ACKR4 | N.A. | 0.15 | 83.4 ± 2.5 | Browns-Forsythe & Welch ANOVA |
| | | CXCR4 | | | 78.2 ± 14.1 | |
| | | CCR9 | | | 111.3 ± 24.1 | |
| | K364A | ACKR4 | N.A. | 0.46 | 70.0 ± 11.5 | Browns-Forsythe & Welch ANOVA |
| | | CXCR4 | | | 74.7 ± 20.0 | |
| | | CCR9 | | | 85.4 ± 8.2 | |
| | K383A | ACKR4 | N.A. | 0.44 | 39.8 ± 23.0 | Browns-Forsythe & Welch ANOVA |
| | | CXCR4 | | | 51.3 ± 7.7 | |
| | | CCR9 | | | 66.0 ± 30.3 | |
| **8a** | WT | N.A. | | N.A. | 100 ± 11.3 | t-test |

| | | | | | |
|---|---|---|---|---|---|
| | KD-GRK3 | | 0.002 | 25.1 ± 7.0 | |
| **8b** | WT | N.A. | N.A. | 100 ± 24.8 | t-test |
| | KD-GRK3 | | 0.24 | 72.5 ± 24.5 | |
| **8c** | WT | N.A. | N.A. | 100 ± 18.0 | t-test |
| | KD-GRK3 | | 0.49 | 90.9 ± 8.7 | |
| **8d** | WT | N.A. | N.A. | 100 ± 25.8 | t-test |
| | KD-GRK3 | | 0.12 | 57.7 ± 26.5 | |
| **S1** | HEK P | N.A. | N.A. | N.A. | Browns-Forsythe & Welch ANOVA |
| | ΔGRK2/3 | | 0.035 | 57.4 ± 14.1 | |
| | ΔGRK5/6 | | 0.049 | 68.7 ± 12.4 | |
| | ΔQ | | 0.012 | 40.5 ± 11.7 | |
| **S2** | WT | N.A. | N.A. | N.A. | Browns-Forsythe & Welch ANOVA |
| | E332A/E333A | | 0.26 | 88.8 ± 12.3 | |
| | D337A/E339A | | 0.27 | 85.2 ± 17.2 | |
| | Proximal | | 0.71 | 95.4 ± 18.4 | |
| | Distal | | 0.058 | 72.4 ± 12.0 | |
| | All | | 0.011 | 43.9 ± 10.0 | |
| **S3** | WT | N.A. | N.A. | N.A. | Browns-Forsythe & Welch ANOVA |
| | E332A/E333A | | 0.044 | 38.0 ± 13.7 | |
| | D337A/E339A | | 0.0077 | 34.0 ± 6.0 | |
| | Proximal | | 0.029 | 24.5 ± 13.5 | |
| | Distal | | 0.0097 | 40.9 ± 6.1 | |
| | All | | 0.0082 | 18.1 ± 7.7 | |
| **S5** | WT | N.A. | N.A. | N.A. | Browns-Forsythe & Welch ANOVA |
| | R226A | | <0.0001 | 70.4 ± 13.6 | |
| | K230A | | 0.0001 | 67.2 ± 16.2 | |
| | R316A | | <0.0001 | 49.4 ± 10.3 | |

| | | | | | |
|---|---|---|---|---|---|
| | K319A | | <0.0001 | 65.1 ± 8.0 | |
| | K345A | | 0.84 | 100.7 ± 10.1 | |
| | K364A | | 0.02 | 86.8 ± 13.6 | |
| | K383A | | 0.93 | 100.8 ± 23.5 | |
| **S6a** | WT | N.A. | N.A. | N.A. | t-test |
| | KD-GRK3 | | 0.58 | 116.8 ± 44.1 | |
| **S6b** | WT | N.A. | N.A. | N.A. | t-test |
| | KD-GRK3 | | 0.65 | 123.3 ± 76.1 | |
| **S6c** | WT | N.A. | N.A. | N.A. | t-test |
| | KD-GRK3 | | 0.65 | 116.7 ± 54.3 | |
| **S6d** | WT | N.A. | N.A. | N.A. | t-test |
| | KD-GRK3 | | 0.56 | 121.8 ± 54.4 | |
| **S7b** | WT | N.A. | N.A. | 100 ± 9.7 | t-test |
| | 10xST/A | | 0.0015 | 36.9 ± 6.9 | |
| **S7c** | WT | N.A. | N.A. | N.A. | t-test |
| | 10xST/A | | 0.049 | 71.1 ± 8.9 | |
